# Bioactive Natural Products Produced by *Streptomyces* from the Microbiome of Cadaveric Fly Larvae

**DOI:** 10.64898/2026.03.12.711486

**Authors:** Shukria Akbar, Rauf Salamzade, Kaetlyn Ryan, Caitlin M. Carlson, Adam J. Schaenzer, Mostafa Zamanian, Lindsay R. Kalan, Tim S. Bugni, Cameron R. Currie

## Abstract

*Streptomyces* are prolific producers of bioactive compounds and increasingly recognized as members of insect microbiomes, yet the microbiome of cadaveric fly larvae remain an overlooked system for discovering metabolically versatile *Streptomyces* species. Here, we conduct targeted bacterial isolations from the microbiome of fly larvae collected from pig cadavers, generating 42 *Streptomyces* isolates of interest, and systematically evaluated their metabolic potential through genomic analysis, antimicrobial screening, biosynthetic gene cluster assessment, untargeted LC-MS/MS metabolomics, and compound purification. The *Streptomyces* isolates spanned nine species, including underrepresented lineages for which we added genomic representatives. *Streptomyces* from carrion fly larvae exhibited broad-spectrum antimicrobial activity and substantial BGC diversity, supported by metabolomic detection of antimycins, surugamides, and macrotetrolides. From a deep phylogenetic lineage, we purified JBIR-68 and Simamycin and demonstrated their potent anthelmintic activity against *Brugia malayi* microfilariae. GNPS molecular networking revealed three additional JBIR-68 analogs, establishing the first taxonomically resolved *Streptomyces* lineage capable of producing these rare metabolites. Our findings position cadaveric fly larvae as a rich ecological reservoir for discovering *Streptomyces* with the potential to produce chemically diverse natural products with biomedical applications.

## Introduction

Members of the genus *Streptomyces* are renowned for producing structurally diverse natural products with antibacterials, antifungals, anthelmintics, immunosuppressants, and anticancer activities (1–3). Although historically viewed as soil-dwelling bacteria (4,5), recent studies highlight their roles as symbionts with plants and insects (6–9), where they protect host plants, social insect colonies, and insect larvae and food resources through antimicrobial production and may also contribute to nutrient acquisition (10–12). The selective ecological pressures of insect lifestyles have driven the evolution of metabolic specialization in their associated *Streptomyces*, making them a promising source of bioactive natural products (13–15).

Among insect-associated habitats, fly larvae inhabiting cadavers represent a dynamic and largely unexplored microbial niche. Cadavers serve as transient, nutrient-rich environments shaped by complex microbial assemblages originating from endogenous decomposition processes and exogenous inputs such as insects and scavengers (16–19). Microbial succession within decomposing cadavers mediates key ecological and chemical processes, generating volatile and nonvolatile metabolites that influence insect behavior, mediate microbial competition, and structure community dynamics (20–25). Despite the detection of Actinobacteria signatures in these systems (26,27), the presence, diversity, and functional potential of *Streptomyces* in cadaveric fly larvae remain largely uncharacterized. Targeting these larvae therefore offers a unique opportunity to uncover hidden *Streptomyces* diversity and gain insight into their ecological and metabolic roles within this niche.

Here, we systematically investigate *Streptomyces* associated with cadaveric fly larvae to assess their phylogenetic diversity and composition, biosynthetic capacity, and metabolite production. Using an integrated framework that combines phylogenomics, antimicrobial screening, biosynthetic gene cluster annotation, untargeted LC–MS/MS metabolomics, and compound purification, we reveal underrepresented *Streptomyces* species with broad antimicrobial activities and diverse biosynthetic repertoires. This approach further led to the identification of a deep lineage that consistently produces the rare geranylated dihydrouridine and uridine molecules (28,29), along with their novel analogs, and to the discovery of their previously unreported anthelmintic activity. Together, these findings establish cadaveric fly larvae as a rich ecological reservoir of metabolically versatile *Streptomyces* and highlight the value of this niche for accessing hidden genomic and chemical diversity with biomedical potential.

## Results and Discussions

### Cadaveric fly larvae harbor *Streptomyces* species

As part of our continuing interest in exploring the microbial and chemical diversity of insects-associated *Streptomyces* (9), we isolated 51 actinobacterial isolates from 20 fly larvae, 10 collected from each of two pig cadavers from a field trip in Hawaii in 2014. Following Illumina sequencing, genome quality was assessed using thresholds of ≥50% completeness and ≤10% contamination and high quality genomes were taxonomically classified using the Genome Taxonomy Database (GTDB) (30).

Genomes of forty-two out of 51 sequenced isolates shared ≥95% Average Nucleotide Identity (ANI) with the genomes of *Streptomyces* genus in GTDB while remaining belonged to non-*Streptomyces* genera, including *Micromonospora*, *Rhodococcus*, and *Actinomadura* (Fig. S1). *Streptomyces* species with ≥3 isolates were considered likely abundant in cadaveric fly larvae system and labeled as species A–F in this study (Fig. 1 & Fig. S2). Among the *Streptomyces*, *Streptomyces* sp. 010548465 (14 isolates), *Streptomyces diastaticus* (9 isolates), *Streptomyces* sp. 001293595 (5 isolates), *Streptomyces* sp. 000772045 (4 isolates), *Streptomyces ardesiacus* (4 isolates), *Streptomyces albidoflavus* (3 isolates), *Streptomyces sennicomposti* (1 isolate), *Streptomyces* sp. 000154905 (1 isolate), and one potentially novel *Streptomyces* sp., were identified (Fig. S2). The high diversity of *Streptomyces* isolates recovered from cadaveric fly larvae underscores the ecological complexity of this niche and prompted their comprehensive evaluation in this study. Species A (*S. albidoflavus*), species B (*S. diastaticus*), and species F (*S. ardesiacus*) showed substantial representation in the GTDB database, suggesting these are well-characterized *Streptomyces*. In contrast, species C (*S.* sp. 010548465), species D (*S.* sp. 001293595), and species E (*S.* sp. 000772045) were underrepresented in GTDB, indicating lesser characterized isolates. Species C had only seven genome entries, including four strains previously isolated from insect hosts. Similarly, despite its deep evolutionary divergence, species D was represented by only four strains, whereas species E had just a single genome entry. Our cadaveric fly larvae study contributes an additional 14 isolates to species C, 5 to species D, and 4 to species E from a single sampling expedition (Fig. 1 & Fig. S2).

**Figure 1.**
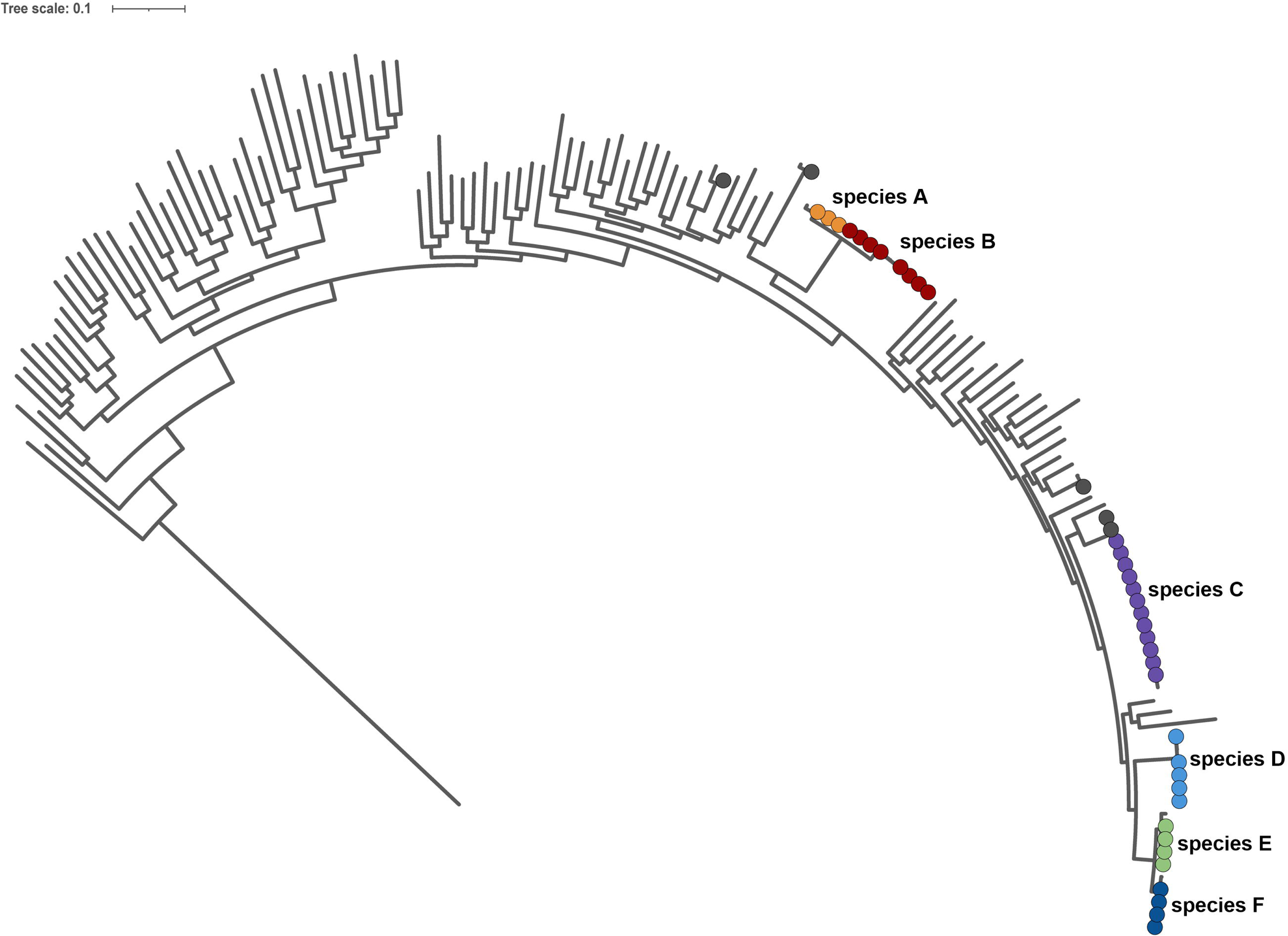
Taxonomic classification of *Streptomyces* isolates recovered from cadaveric fly larvae. A genome-based phylogeny was constructed using the 42 *Streptomyces* isolates obtained in this study, six GTDB R214 reference genomes representing the recovered species, and an additional set of 100 diverse *Streptomyces* genomes to provide broader taxonomic context. The phylogeny was generated from GTDB-Tk concatenated marker genes and visualized in iTOL. Isolates recovered from cadaveric fly larvae are labeled as “SID” and highlighted with colored nodes, each corresponding to one of the six species-level groups (A–F) identified in this study. The scale bar represents substitutions per site.

The scarcity of soil-derived representatives for these three species suggests niche specificity and a likely association with insect hosts. Overall, this highlights cadaveric fly larvae system as an overlooked reservoir for uncovering less abundant *Streptomyces* lineages and accessing underexplored genomic and functional diversity.

### Isolated *Streptomyces* exhibit antagonistic activities towards pathogens

Given that *Streptomyces* are known components of the insect microbiome and often act as beneficial symbionts that protect insect colonies, larvae, and food resources from microbial antagonists (9–11), we sought to evaluate the antimicrobial capacity of the *Streptomyces* isolates recovered from cadaveric fly larvae. Using a co-culture competition assay against a panel of Gram-positive and Gram-negative bacteria as well as fungal pathogens, we compared the antagonistic activities of these isolates (31).

All isolates demonstrated inhibitory activity against the tested pathogens, with members of species A and B particularly showing complete inhibition of fungal pathogens (Fig. 2). Isolates of species A and B also displayed comparatively good Gram-positive antibacterial activity. Among the isolates of species C, all except SID7959 and SID8458 exhibited substantial inhibition, and several completely inhibited MRSA and *Aspergillus flavus*. Species D isolates displayed broad-spectrum activity, with SID7919 and SID9885 fully inhibiting fungal pathogens and Gram-negative bacteria, including *Acinetobacter* spp. Similarly, species E isolates produced moderate to strong inhibition, with SID10362 fully suppressing *Pseudomonas aeruginosa*. Species F isolates demonstrated variable activity, predominantly targeting *Aspergillus* and Gram-positive pathogens.

**Figure 2.**
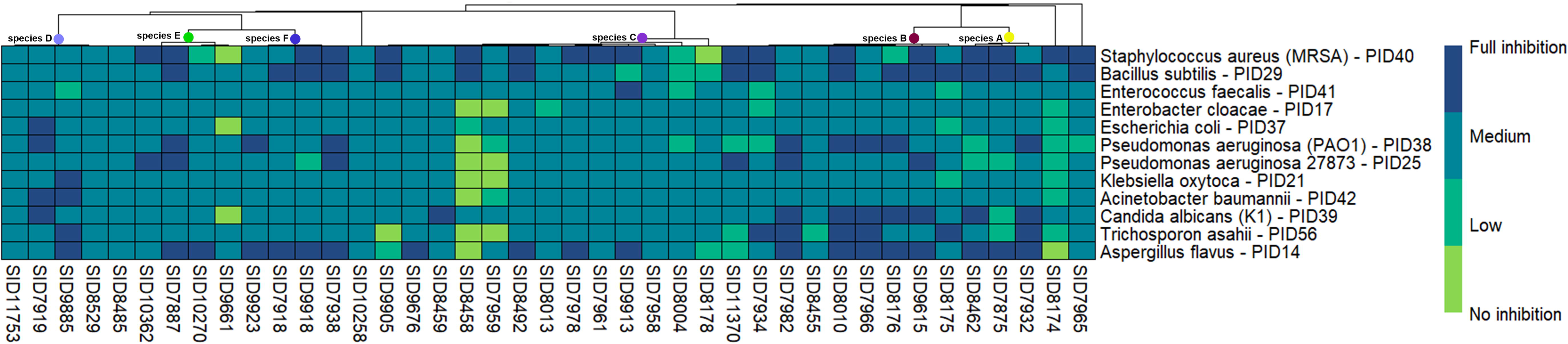
Inhibitory activity of *Streptomyces* isolates against bacterial and fungal pathogens. Antimicrobial activity of the *Streptomyces* isolates was evaluated using a co-culture competition assay against a panel of clinically relevant bacterial and fungal pathogens. Colored nodes above the heatmap denote the species-level groups (A–F) corresponding to each isolate. Pathogen inhibition was scored semi-quantitatively, where neon indicates no inhibition, green indicates slight inhibition, cyan indicates a clear zone of inhibition, and blue represents complete inhibition. The heatmap was generated in R (version 4.4.3) using the pheatmap package.

Despite generally consistent antimicrobial patterns, variation among taxonomically related isolates suggests inherent functional disparities (32,33). The spectrum and strength of the observed antagonistic activities suggest that *Streptomyces* from cadaveric fly larvae possess broad and robust antimicrobial capacities and likely play an active role in shaping microbial interactions within this niche.

### *Streptomyces* from Cadaveric fly larvae demonstrate enriched biosynthetic capacity

To investigate the biosynthetic potential of the cadaveric fly larvae *Streptomyces* isolates, we annotated biosynthetic gene clusters (BGCs) in their genomes using antiSMASH (34) and BiG-SCAPE (35).

This analysis revealed substantial heterogeneity in BGC distribution across species, with only a few conserved clusters (Fig. 3). Multiple isolates of species A and species B contained BGCs for polycyclic tetramate macrolactam (BGC0001043; BGC0002509; BGC0000996) (36), antimycins (BGC0000958; BGC0001216; BGC0001455) (37), and candicidin (BGC0000034) (38), the compounds known for broad biological activities, including antifungal activities (Fig. 3 & Fig. 4A). One isolate of species B harbored a BGC for an anti-influenza compound, violapyrone B (BGC0001905) (39). Species C isolates carried the weishanmycin BGC (BGC0001823) (40), while Species D encoded collinomycin (BGC0000266) (41), responsible for synthesizing compounds with cytotoxic and antibacterial activity, respectively. A cluster for Gram-positive bacterial active curamycin (BGC0000215) (42) was present in the isolates of species D, E, and F. Species E and F also shared the albaflavinone BGC (BGC0000660), responsible for producing a compound active against *Bacillus subtilis* (43). Species E additionally harbored species-specific clusters for the cytotoxic compound ashimide (BGC0002288) (44), the Gram-positive–active antibiotic α-lipomycin (BGC0001003) (45), and the highly bioactive undecylprodigiosin (BGC0001063) (46), while two isolates of species F encoded venemycin (BGC0001819) (47). These patterns highlight both shared and species-restricted biosynthetic capacities across lineages.

**Figure 3.**
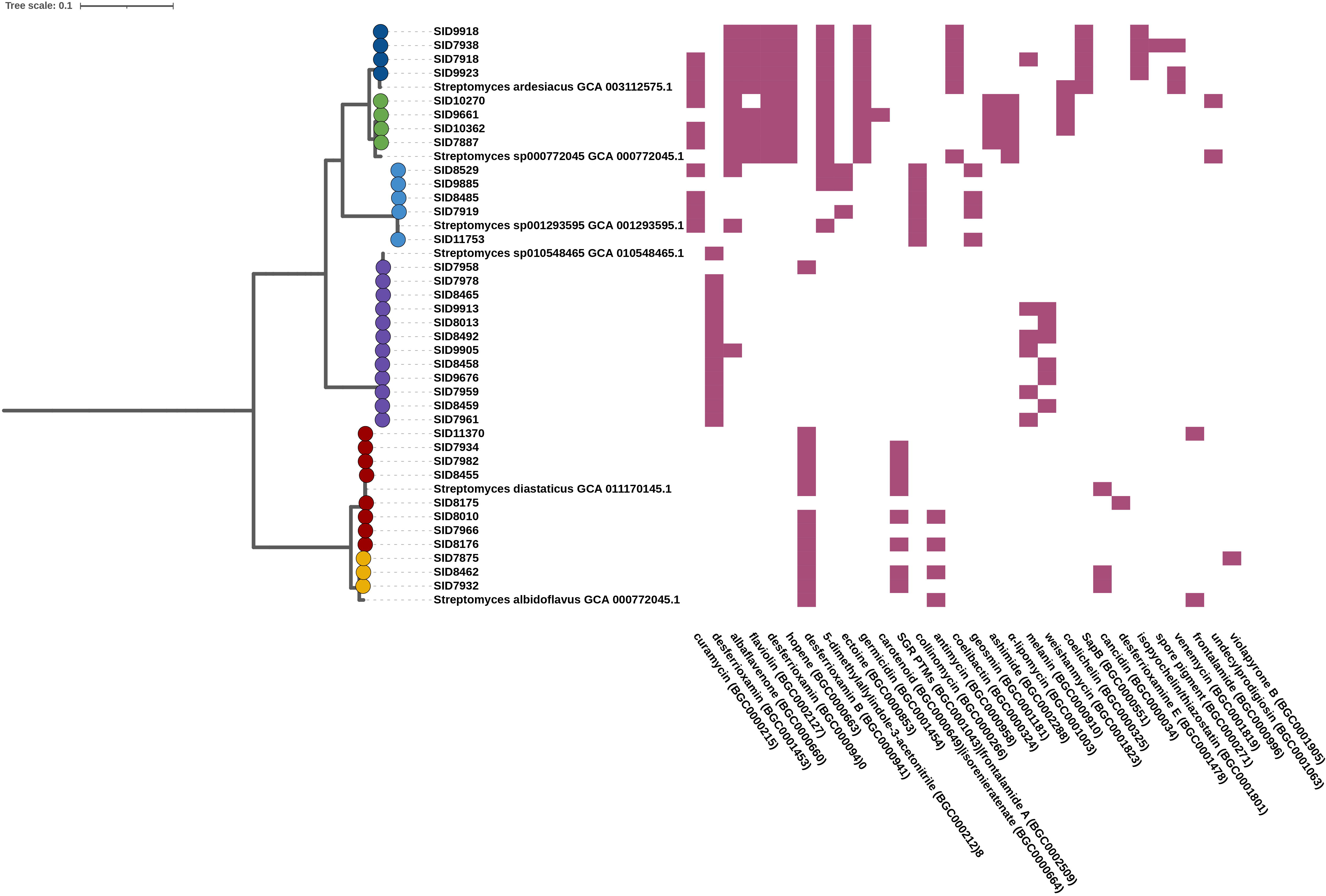
Distribution of characterized biosynthetic gene clusters across *Streptomyces* isolates and their reference genomes. Characterized BGCs detected in the genomes of the isolates obtained in this study (colored nodes) and species-representative genomes included in the phylogeny are shown. Each column represents a gene cluster family containing at least one MIBiG-characterized BGC, and each filled cell denotes the presence of that BGC in a given genome. The phylogeny was generated from GTDB-Tk concatenated marker genes, and both the tree and the BGC presence–absence matrix were visualized using iTOL. The scale bar represents substitutions per site.

**Figure 4.**
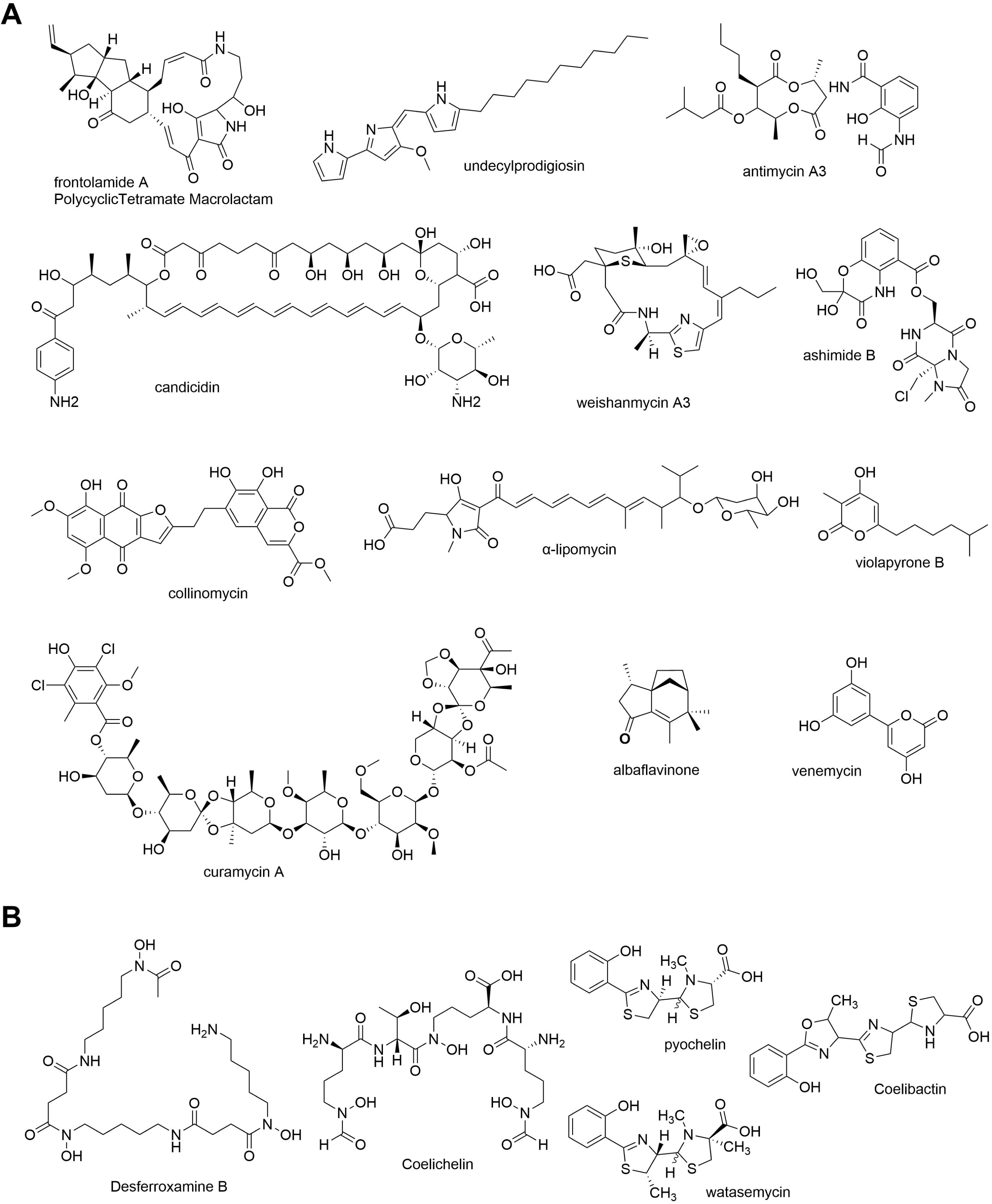
Metabolites associated with biosynthetic gene clusters detected in *Streptomyces* isolates from cadaveric fly larvae. (A) Structures of bioactive specialized metabolites corresponding to BGCs identified across the genomes of the *Streptomyces* isolates in this study (B) Structures of siderophores associated with conserved BGCs detected in multiple isolates.

Alongside bioactive metabolites, BGCs for diverse metals acquiring siderophore were present across isolates. Desferrioxamine gene clusters (BGC0001453; BGC0000940; BGC0000941; BGC0001478) (48) occurred in all species except species D, while coelichelin (BGC0000325) and coelibactin (BGC0000324) (49) were restricted to species E and species F, respectively (Fig. 3 & Fig. 4B). Species F also encoded pyochelin-type siderophores such as watasemycin (BGC0001801) (50). The presence of chemically distinct siderophore pathways indicates functional redundancy in metal acquisition, consistent with strong resource competition in the cadaveric larvae niche. Collectively, the isolates exhibit broad and varied biosynthetic potential, suggesting a metabolically versatile community adapted to competitive environments.

### *Streptomyces* from cadaveric fly larvae produce bioactive metabolites

To more fully assess the metabolic potential of the cadaveric fly larvae *Streptomyces* isolates, we performed untargeted LC-MS/MS metabolomic profiling on methanolic crude extracts and analyzed the data using GNPS (51,52) and Bruker ProfileAnalysis (Bruker Daltonics, 2018).

GNPS-rendered molecular networks detected antimycins in species A and B (53), surugamides (54) in species B, and macrotetrolides (55) in species C (Fig. 5 & Fig. S3). Notably, metabolite presence and intensity varied among the isolates of a species. For species A, only one of three isolates (SID7875) produced antimycin A1 (*m/z* 549.276) and A2 (*m/z* 535.267), whereas antimycin A1, A2 and A3 (*m/z* 521.244) were detected in only five of the eight isolates of species B (Fig. S4). Similarly, all species B isolates except SID8455 and SID8176 consistently produced surugamides A (*m/z* 912.624), B (*m/z* 898.609), G (*m/z* 884.599), and H (m/z 870.575) with surugamide A highest and H lowest in abundance (Fig. 5 & Fig. S5-S7). Ten of the fourteen isolates of species C produced cyclic macrotetrolides, including monactins (*m/z* 768.486), dinactin (*m/z* 782.501), trinactin (*m/z* 796.516), tetranactin (*m/z* 810.532), and nonactin (*m/z* 754.489) with dinactin showing the highest ion abundance (Fig. 5 & Fig. S8). Bonactin (*m/z* 423.234) (37), an acyclic polyether, was detected alongside these metabolites (Fig. 5 & Fig. S9). Although corresponding BGCs for macrotetrolides and surugamides were not detected in Illumina-assembled genomes, their metabolomic detection suggests additional biosynthetic traits not captured genomically.

**Figure 5.**
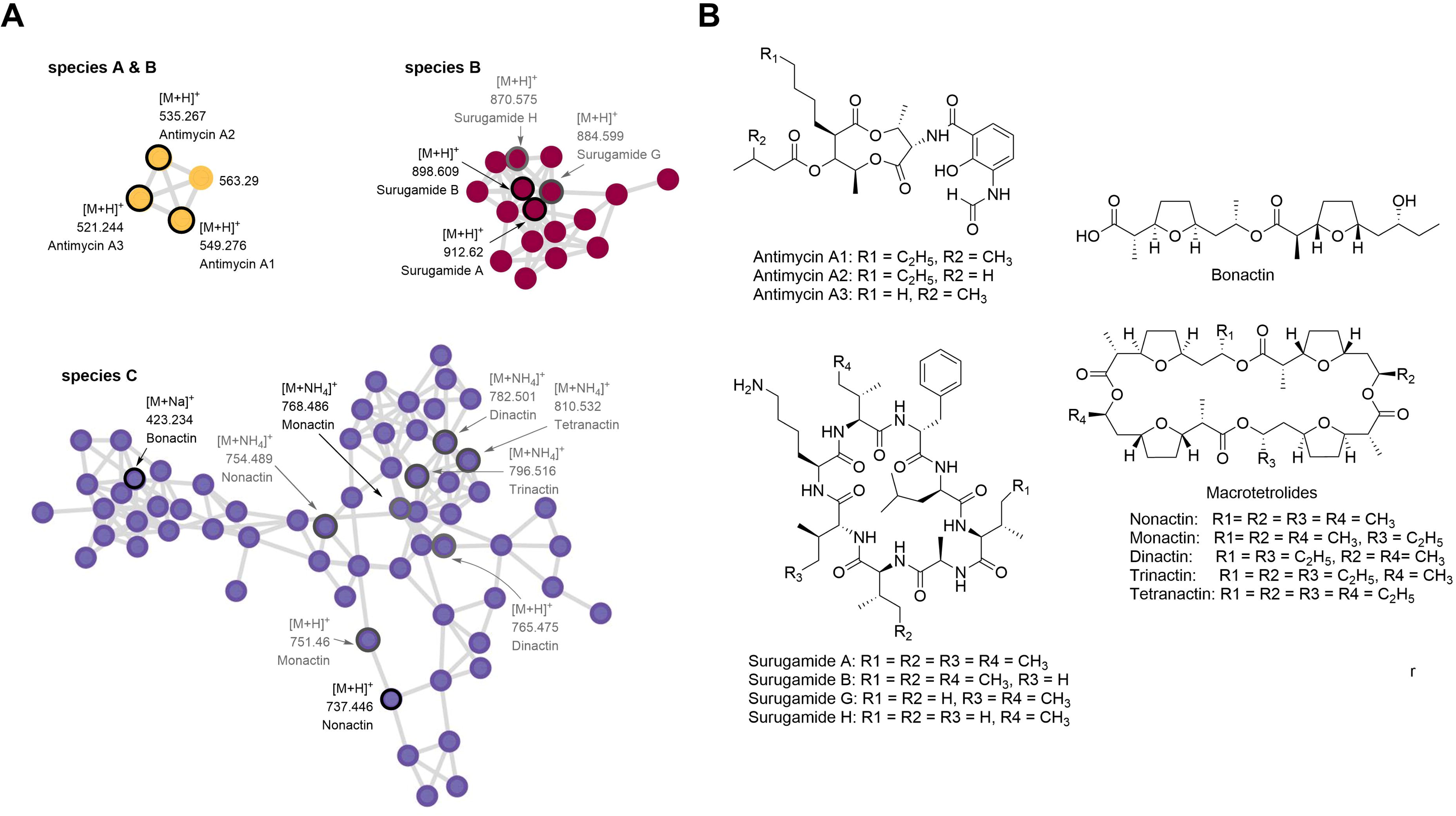
Metabolites detected in crude extracts of *Streptomyces* isolates recovered from cadaveric fly larvae. (A) GNPS-rendered molecular networks highlighting mass features corresponding to antimycins detected in isolates of species A and B, surugamides detected in species B, and macrotetrolides detected in species C. Each node represents a mass feature; nodes with black labels indicate GNPS library matches, while nodes with grey labels denote features annotated based on literature references. (B) Chemical structures of the metabolites corresponding to the detected mass features.

No previously known metabolites were detected in crude extracts from the isolates of species D, E, or F, including those predicted from their BGCs. This absence likely reflects the lack of matching compounds in GNPS libraries or potential novelty in their metabolomes. Variation in metabolite profiles across isolates underscores functional diversity among the related isolates (56). These findings highlight the value of sampling multiple isolates to capture chemical diversity and demonstrate that cadaveric fly larvae represent a rich, underexplored niche for metabolically diverse *Streptomyces* with significant biomedical potential.

### A *Streptomyces* isolate from cadaveric fly larvae produces geranylated dihydrouridine and uridine molecules

The deep phylogenetic divergence of species D (Fig. 1), combined with limited biosynthetic and metabolomic information of its isolates, prompted further investigation. Bioactivity data identified SID9885 and SID7919 as promising isolates (Fig. 2). HPLC purification and chemical analysis of the SID9885 crude extract led to the characterization of 5’’-O-geranyl dihydrouridine (1) and a semipurified compound, 5’’-O-geranyl uridine (2) (Fig. 6).

**Figure 6.**
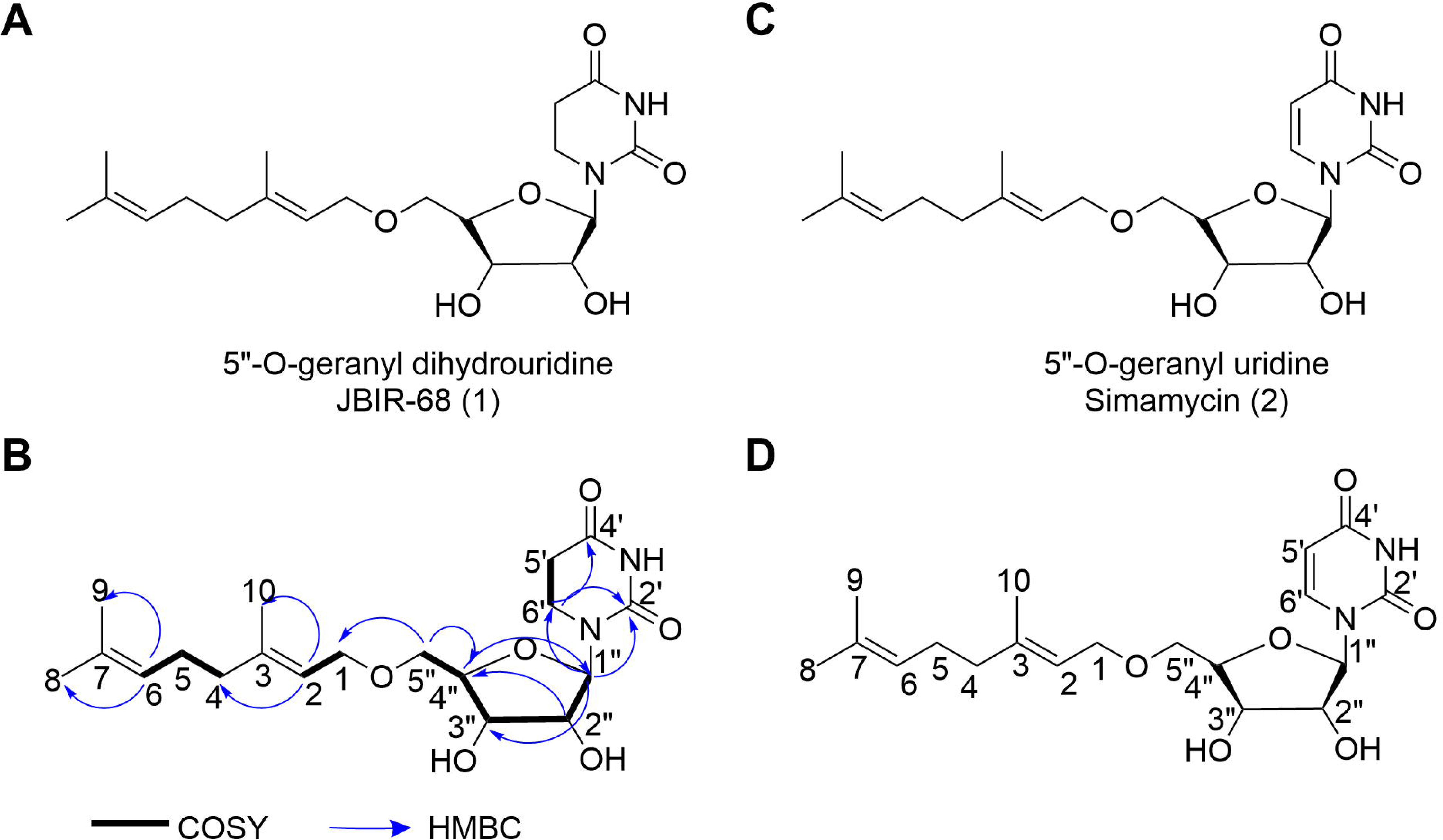
Structural characterization of JBIR-68 and Simamycin purified from an isolate of species. **D.** (A) Chemical structure of JBIR-68. (B) Key COSY and HMBC correlations used to establish the structure of JBIR-68. (C) Chemical structure of Simamycin (5ʺ-O-geranyl uridine). (D) Simamycin structure with atom numbering used for comparative structural analysis.

Compound 1 was purified as a colorless substance with [M+Na]^+^ at 405.199, consistent with C_19_H_30_N_2_O_6_ and supported by 1H and 13C NMR data (Fig. S10-S14; Tab. S1). Analysis of 1D and 2D NMR data confirmed a geranyl and a dihydrouridine moiety (Fig. 6B). Full characterizing of the molecule confirmed the structure as 5’’-O-geranyl dihydrouridine. Comparison with published data established the compound as JBIR-68 (28). A semipurified fraction containing compound 2 showed [M+Na]^+^ at 403.183 for the molecular formula C_19_H_28_N_2_O_6_ (Fig. S14). Its 2.01 Da lower mass than JBIR-68 and the presence of two olefinic protons at 5’ (8.06 ppm, 1H, d, J=8.14) and 6’ (5.61 ppm, 1H, d, J=8.14) positions (data not shown), instead of methylene signals, supported its identification as 5’’-O-geranyl uridine, previously described as Simamycin (Fig. 6C & D) (29).

Both JBIR-68 and Simamycin are likely biosynthesized through a single enzymatic geranylation of dihydrouridine/uridine, explaining the absence of a corresponding BGC in antiSMASH predictions. Notably, JBIR-68 and Simamycin have been reported only once previously (28,29). Our findings provide new evidence for their production and highlight the unique biosynthetic potential of cadaveric fly larvae associated *Streptomyces*.

### JBIR-68 and Simamycin demonstrate anthelmintic activity

JBIR-68 and Simamycin were previously reported to exhibit anti-influenza activity and to induce preadipocyte differentiation, respectively (28,29). The absence of documented antimicrobial activity for these molecules aligns with our screening results. However, given that they possess a geranyl moiety, a terpene known for its antinematodal activity (57–60), we evaluated the effects of JBIR-68 and Simamycin against *Brugia malayi* microfilariae, the parasitic nematode responsible for lymphatic filariasis in humans (61).

Both compounds significantly reduced worm motility at 100 µg/mL after 24 hours, with complete inhibition by 48 hours (Fig. 7A). Parasite viability was also impaired at this concentration indicating that geranylated uridines possess notable anthelmintic activity. This represents the first report demonstrating the anthelmintic potential of geranylated uridine and dihydrouridine derivatives. As species D consists of four additional isolates besides SID9885, we assessed their semicrude fractions against *B. malayi*. Each fraction inhibited parasite motility and viability at 50 µg/mL with activity slightly exceeding that of the purified compounds, suggesting likely contributions from additional analogs (Fig. 7B). Overall, all species D isolates exhibit anthelmintic potential, highlighting a promising chemical space and offering insight into how this lineage and geranylated nucleoside molecules may mediate interactions within cadaveric systems.

**Figure 7.**
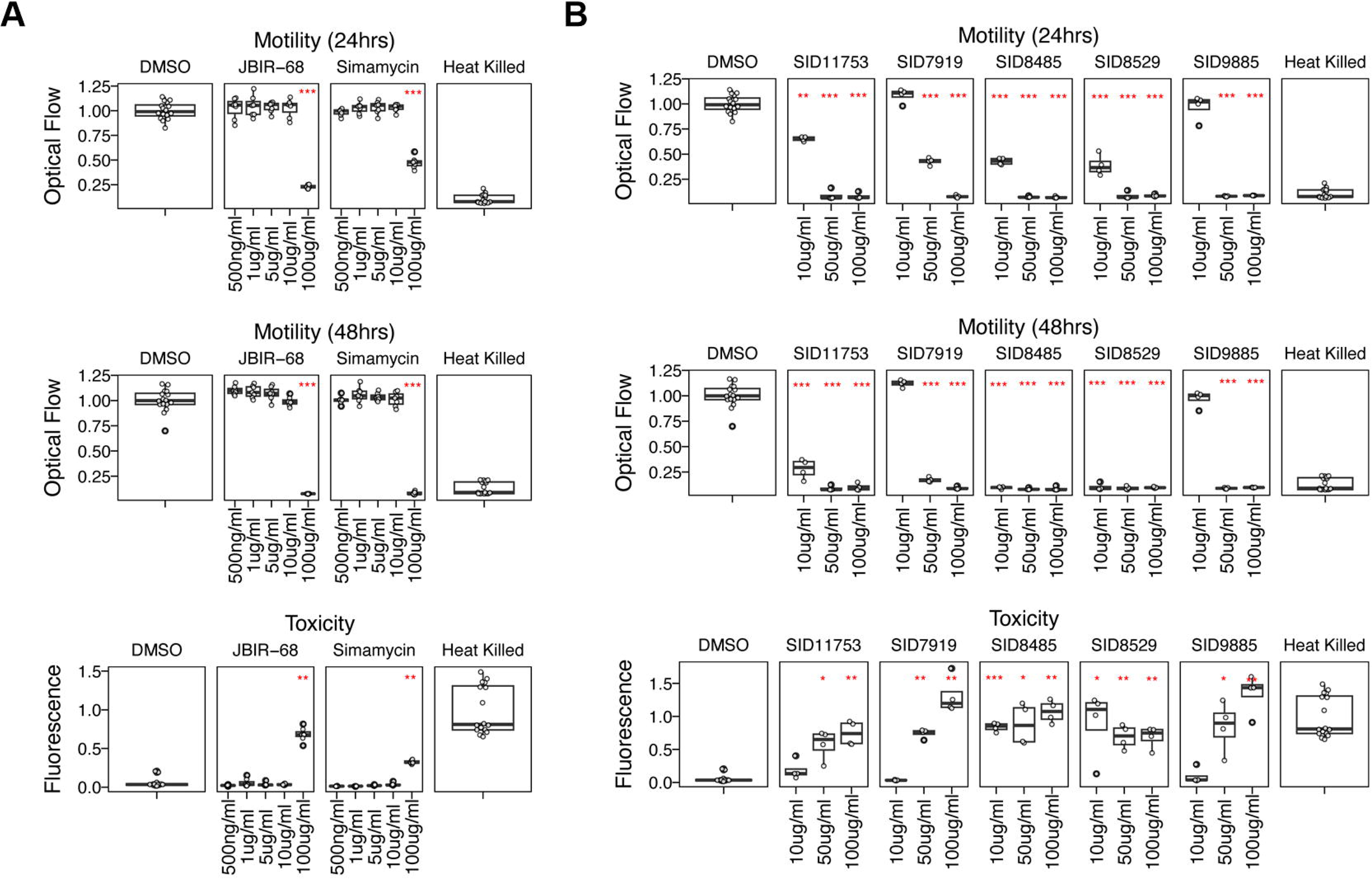
Anthelmintic activity of JBIR-68, Simamycin, and semicrude extracts from species D isolates against *Brugia malayi* microfilariae. (A) Anthelmintic activity of purified JBIR-68 and Simamycin against *Brugia malayi* microfilaria. (B) Anthelmintic activity of semicrude extracts from species D isolates. Worm motility was quantified at 24 and 48 hours post-treatment using the ImageXpress Nano, and worm viability was assessed by fluorescence measurement using the CellTox Green Cytotoxicity Assay. Data were normalized and visualized in R (version 4.4.3) using the tidyverse package. Heat-killed worms served as the positive control, and DMSO as the negative control.

### Identification of a *Streptomyces* lineage producing JBIR-68 and its analogs

To determine the presence of JBIR-68, Simamycin and their analogs in semicrude fractions of species D isolates, we performed untargeted LCMS/MS metabolomics and GNPS molecular networking (52).

JBIR-68 and Simamycin were detected in the semicrude fractions of all five isolates. Although Simamycin was not detected in a molecular network, the extracted ion chromatogram (EIC) confirmed its presence in all the fractions (Fig. S15). JBIR-68 along with its analogs was detected in a distinct molecular subnetwork within a larger GNPS rendered molecular network (Fig. 8B). Mass differences between connected nodes, where each node represents a distinct mass feature, indicates presence of three additional JBIR-68 analogs (Fig. 8B).

**Figure 8.**
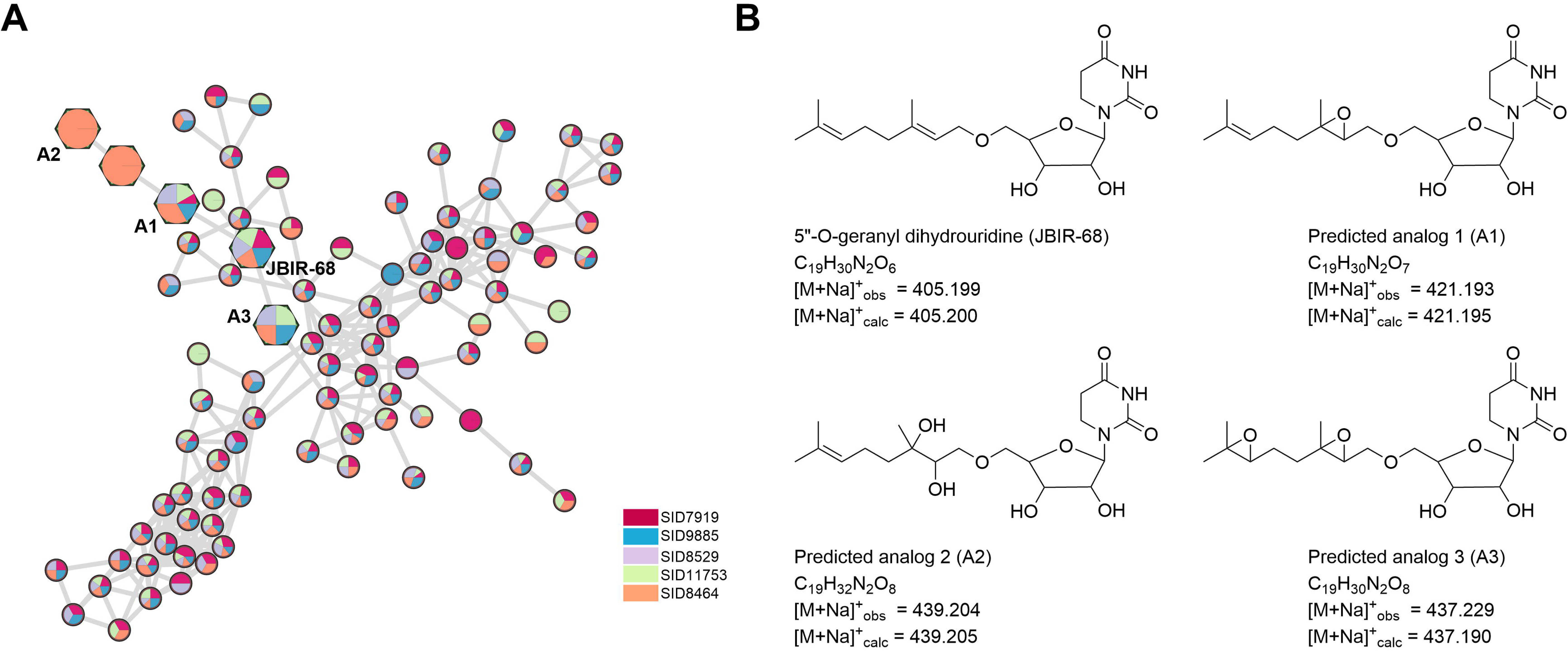
JBIR-68 and predicted structural analogs detected through GNPS molecular networking. (A) GNPS-rendered molecular network highlighting the subnetwork containing the mass feature for JBIR-68 and its predicted analogs. Diamond-shaped nodes represent features belonging to this subnetwork, and pie-chart coloring indicates the proportional contribution of each isolate to the corresponding mass feature. (B) Chemical structure of JBIR-68 (5ʺ-O-geranyl dihydrouridine) and the predicted analogs inferred from observed mass differences, with calculated and observed [M+Na]⁺ values shown for each compound.

JBIR-68 and predicted analog 1 (A1) were detected in all isolates. Predicted analog 3 (A3) was present in all except SID9885, whereas predicted analog 2 (A2) was unique to SID8485 (Fig. 8). We confirmed the presence of these detected mass features through the extracted ion chromatogram (EIC) in the corresponding strains (Fig. S16). The presence of these additional analogs likely contributed to the slightly enhanced anthelmintic activity observed in semicrude fractions (Fig. 8B). Although JBIR-68 and Simamycin have been previously isolated from *Streptomyces*, the taxonomic identity of their producing strains was never established (28,29). Our findings provide the first taxonomically resolved *Streptomyces* lineage known to produce these compounds, along with three detected novel analogs, expanding the known chemical capacity of this underexplored species. This study also underscore the value of sampling the competitive cadaveric fly larvae niches to capture multiple related isolates and assess chemical diversity.

## Conclusion

Our integrative genomic, metabolomic, and functional analysis reveals that cadaveric fly larvae represent a rich ecological reservoir of phylogenetically and metabolically diverse *Streptomyces*. By expanding genomic representation for several underexplored *Streptomyces* species and identifying the first taxonomically resolved producers of JBIR-68, Simamycin, and related analogs, we uncover hidden chemical capacity within this niche. The demonstrated anthelmintic and antimicrobial activities highlight the ecological relevance of these metabolites and underscore the potential of cadaveric systems for natural product discovery. Together, these findings illustrate the value of environmentally guided sampling for accessing untapped microbial diversity and biosynthetic innovation.

## Materials and Methods

### Collection of fly larvae

Fly larvae-associated strains were obtained from the larvae collected from two pig cadavers from a field trip in Hawaii in 2014. The larvae were collected using sterile forceps and deposited into a pre-sterilized, pre-barcoded container. Collections were focused heavily on larvae not in direct contact with soil to avoid the possibility of soil contamination in subsequent bacterial isolation. The collected larvae were stored at 4 °C

### Processing and bacterial isolation

Bacterial isolation was performed according to Chevrette et al (9). Briefly, larvae were transferred into a 1.5 mL microcentrifuge tube, and phosphate-buffered saline (PBS) was added at a volume of 125*x, where x represents the number of agar plates used for inoculation. Samples were gently vortexed at 50% speed for 10 sec to dislodge surface-associated microbes. A 100 μL aliquot of the resulting suspension was subsequently spread onto various isolation media. To selectively culture Actinobacteria, humic acid agar (HV) (62) and selective chitin media (63) supplemented with 20 mL/1L nystatin, and 10mL/1L cycloheximide were employed. Plates were incubated aerobically at 28 °C and monitored for colony development at 14, 30, and 90 days. Colonies exhibiting characteristic Actinobacterial morphology were assigned unique strain identification (SID) numbers and selected for subsequent DNA extraction, and antimicrobial activity assays.

### DNA sequencing and assembly

Based on distinct morphology, bacterial Isolates were selected for whole genome sequencing. Here, the DNA extraction method for one sample is explained. Bacterial culture was grown on a rich medium supplemented with 0.5% glycine. Cells were harvested by centrifugation and washed with 10.3% sucrose. The pellet was resuspended in a lysozyme solution (3 mg/mL lysozyme, Sigma) in 0.3 M sucrose, 25 mM Tris (pH 8), and 25 mM EDTA (pH 8) and incubated at 37 °C for 30 min. Following lysozyme treatment, Proteinase K (Thermo Fisher; 20 mg/mL) was added, and the mixture was incubated for an additional 15 min at 42 °C. Cells were lysed by adding 2% SDS and gently shaken for 5 minutes until complete lysis was achieved. The lysate was then extracted with neutral phenol and chloroform, with gentle shaking until a uniform white phase was observed. After centrifugation, the aqueous phase was transferred to a new tube containing 3 M sodium acetate (pH 6) and isopropanol, and DNA was precipitated by gentle mixing. The DNA pellet was recovered by centrifugation, and the supernatant was discarded. The pellet was resuspended in TE buffer with 0.2 mg/mL RNase A and incubated for 15 min at 28 °C. Then, NaCl (5 M) and CTAB/NaCl solution were added, and the tubes were incubated for 10 min at 55 °C before being cooled to 28 °C. Finally, chloroform was added to the tube, and the mixture was gently shaken and centrifuged at 28 °C for 10 minutes. The aqueous phase was transferred to a fresh tube and re-extracted with phenol and chloroform, followed by a final extraction with chloroform. DNA was then precipitated with 3 M sodium acetate (pH 6) and isopropanol. The resulting DNA pellet was washed with 70% ethanol and resuspended in water. DNA concentration and purity were assessed. Genomic DNA libraries for Illumina MiSeq 2× 300bp paired-end sequencing were prepared by the University of Wisconsin-Madison Biotechnology Center (TruSeq). Reads were corrected with MUSKET v1.153 (64), paired-ends were merged with FLASH v1.2.754 (65), and assembled with SPAdes v3.11.055 (66). All sequenced genomes were deposited in NCBI database under BioProject PRJNA1428808.

### Taxonomic classification, phylogeny construction, visualization

Isolates which were whole-genome sequenced for this study were taxonomically classified using GTDB-Tk (v2.3.2) (67) with GTDB R214 (30).

To provide evolutionary context for *Streptomyces* genomes from this study, 100 diverse *Streptomyces* genomes were determined. Briefly, 1,555 strain-distinct *Streptomyces* genomes from GTDB R214 (30) following average nucleotide identity-based dereplication using skDER (68,69) were downloaded and gene calling was performed using pyrodigal (v3.6.3) (70). Afterwards, GToTree (v1.8.8) (71) was used to annotate largely single-copy core proteins for Actinomycetota and to construct an approximate maximum-likelihood phylogeny using FastTree 2 (v2.1.11) (72). A hundred distinct *Streptomyces* genomes were selected from the phylogeny using treemmer (v0.3) (73).

For construction of the phylogeny shown in Figure 1 & 2, GToTree and FastTree 2 were used on a set of 148 *Streptomyces* genomes with one *Spirillospora geliboluensis* (SID9922) genome also included for use as an outgroup for rooting. The *Spirillospora* genome was removed for visualization. The 148 *Streptomyces* genomes included the set of 100 distinct *Streptomyces* representative genomes from treemmer dereplication, the 42 *Streptomyces* genomes from this study, and six representative genomes from GTDB R214 for species represented by the genomes from this study. One *Streptomyces diastaticus* genome (SID9615) was automatically filtered from inclusion in the phylogeny due to featuring too few single-copy core proteins. This isolate was the only one determined as likely contained based on CheckM2 (v1.0.1) (74) analysis. To assess strain similarity amongst the 148 genomes in the phylogeny, skani (v0.2.2) (69) was used to determine whether pairs of genomes exhibited ≥80% average nucleotide identity and ≥50% bi-directional coverage to each other. Phylogenetic visualizations were performed using iTol (75).

### Antimicrobial assay

High throughput co-culture assay according to Temkin et al (31) was utilized to test the antimicrobial potential of the isolates. Isolated bacteria were inoculated onto one side of each well in a 12-well plate containing 3 mL of yeast peptone mannitol (YPM) agar (2 g yeast extract, 2 g peptone, 4 g mannitol, 15 g agar, 1 L water). Plates were incubated at 28 °C for 5 days to allow for growth of the actinobacterial isolates prior to pathogen introduction. For fungal pathogens, spore suspensions were prepared by diluting stock solutions 1:10 in sterile water. Bacteria and yeast pathogens were cultured in 3 mL of broth of Luria-Bertani (LB) or yeast peptone dextrose (YPD), respectively. Cultures were then diluted 1:10, and 3 μL of each diluted pathogen culture was inoculated onto the side of the well opposite the *Streptomyces* inoculum. Plates were incubated at 28 °C for an additional 7 days. Pathogen inhibition was assessed on a semi-quantitative scale from 0 to 3, where 0 indicated no inhibition, 1 indicated slight inhibition, 2 indicated a clear zone of inhibition, and 3 indicated complete inhibition. A heatmap representing the pathogen inhibition scores was generated in R (version 4.4.3) using pheatmap package (76).

### Annotation and determination of characterized and novel BGCs

To annotate for the presence of BGCs, we applied the software antiSMASH (v8.0.2) (34) with parameters: “--taxon bacteria --genefinding-tool prodigal --fullhmmer --asf --cb-general --cb-subclusters --cb-knownclusters --cc-mibig --rre --pfam2go” on each genome from this study and species-representative genomes included in phylogenies. Gene cluster families (GCFs) and the presence of characterized BGCs in the MIBiG database (v3.1) (77) for the same set of genomes were determined using BiG-SCAPE (v1.1.5) (35). For running BiG-SCAPE, we applied the options “--mibig --include_singletons --mix”. BiG-SCAPE mixed clustering results were processed to determine GCFs which featured both a BGC region from a study-specific genome and a characterized MIBiG BGC. Some GCFs were found to feature multiple reference BGCs from MIBiG.

### Acquisition of LC-MS/MS data

*Streptomyces* isolates were cultured on International *Streptomyces* Project-2 (ISP-2) agar medium (4 g yeast extract, 4 g dextrose, 10 g malt extract, 15 g agar, 1 L water) until sporulated. Two 8mm diameter agar cores were sampled directly from the sporulated bacterial plates, extracted with 2 mL of MeOH, and the resulting extracts were vacuum dried. For LCMS/MS sample preparation, dried samples were dissolved in 500 μL of 10% MeOH and loaded onto the solid phase extraction (SPE) column and eluted with MeOH into LC/MS certified vials. For LCMS/MS data acquisition, we utilized Bruker maXis II Ultra-High-Resolution LC-QTOF mass spectrometer (Bruker Scientific LLC., Billerica, MA, USA) coupled to a Waters Acquity H-Class UPLC system (Waters, Milford, MA, USA) and operated by the Bruker Hystar 3.2 software. Chromatographic gradients were performed with a mixture of methanol and water (containing 0.1% formic acid) on an RP C-18 column (Phenomenex Kinetex 2.6 μ m, 2.1 mm × 100 mm; Phenomenex, Torrance, CA, USA) at 0.3 mL/min. The method was as follows: 0-1 min (10% MeOH in H2O), 1–12 min (10%–97% MeOH in H2O), and 12–15.5 min (97% MeOH in H2O). A mass range of m/z 50–1550 was measured in positive ESI mode for all spectra.

### Metabolomics studies

Generated LCMS/MS raw data were converted to open source .mzML format using MS-Convert (78). Metabolite dereplication was performed using the Global Natural Products Social Molecular Networking (GNPS) platform (51) (http://gnps.ucsd.edu), utilizing both the GNPS spectral library search and molecular networking features. LCMS/MS data for this analysis were also deposited in the MassIVE public repository (MSV000100887 & MSV000100884). When a metabolite was detected in one or more isolates, Bruker ProfileAnalysis software (Bruker Daltonics, 2018) was used to determine its presence of the same mass feature (identical *m/z* and retention time) in other isolates.

### Compound Purification

Streptomyces strain SID9885 was cultured in 10 L of yeast peptone mannitol (YPM) broth (2 g/L yeast extract, 2 g/L peptone, 4 g/L mannitol) supplemented with 70 g of HP20 Diaion resin (Sigma-Aldrich). Cultivation was carried out in Fernbach flasks at 28 °C at 200 rpm for 14 days. The HP20 resin was filtered and soaked with acetone for an hour for extraction. After fermentation, the HP20 resin was separated by filtration and extracted with acetone for 1 hour. The acetone extract was filtered, concentrated under reduced pressure, and the resulting crude extract was sequentially partitioned with hexane and chloroform. The chloroform-partition was further purified by size-exclusion chromatography using an LH20 column (GE Healthcare). Final purification of two metabolites was achieved by reversed-phase high-performance liquid chromatography (HPLC) on a C18 semipreparative column (Phenomenex Luna C18(2), 5 μm, 250 × 10 mm). The mobile phase consisted of acetonitrile (ACN) and water containing 0.1% acetic acid, with a flow rate of 3.2 mL/min. The gradient program was as follows: 1–3 min, isocratic at 80% MeOH–H₂O; 3–20 min, linear gradient from 80% to 100% MeOH; 20–22 min, isocratic at 100% MeOH; 22–22.5 min, linear gradient from 100% to 80% MeOH; and 22.5–27.5 min, isocratic at 80% MeOH–H₂O.

### Structure elucidation

The structure of compound JBIR-68 was elucidated using a series of 2D NMR experiments, including HSQC, COSY, and HMBC spectra. High-resolution electrospray ionization mass spectrometry (HRESIMS) was employed to confirm the molecular formulas of both compounds JBIR-68 and Simamycin. The structure of Simamycin was supported by 1D proton NMR data in combination with HRESIMS.

### Anthelmintic activity

*Brugia malayi* microfilariae were obtained from the NIH/NIAID Filariasis Research Reagent Resource Center (FR3), and morphological voucher specimens are stored at the Harold W. Manter Laboratory of Parasitology, University of Nebraska (accession numbers P2021–2032) (79). Parasites were maintained in RPMI 1640 medium supplemented with 0.1 mg/mL penicillin/streptomycin at 37 °C under 5% CO₂.

Purified compounds and fractions were resuspended in dimethyl sulfoxide (DMSO) and diluted to 100x the final assay concentration. Microfilariae were filtered using a PD-10 column to eliminate cellular debris, embryos, and dead worms. For each assay, 1 μL of the diluted DMSO sample was added to individual wells of a 96-well plate, followed by 100 μL of tittered mf in aliquots of 1000 mf per well. Heat-killed positive controls were prepared by incubating mf at 60 °C for 1 hour prior to plating. Plates were imaged for motility at 24 and 48 hours post-treatment using the ImageXpress Nano (Molecular Devices), with environmental controls set to 37 °C and 5% CO₂. Motility was recorded at 4x magnification with 10-frame acquisitions per well. After the 48-hour time point, viability was assessed using the CellTox Green Cytotoxicity Assay (Promega). Staining was performed by incubating plates with the CellTox working solution for 30 minutes at 37 °C. Plates were then washed twice with M9 buffer using the AquaMax 2000 plate washer (Molecular Devices), with centrifugation between washes to prevent microfilariae loss. Final viability imaging was performed at 4x magnification, capturing all four quadrants of each well. Motility and viability images were analyzed using the motility and mf_celltox modules of the *wrmXpress* software (80). Quantitative software output was normalized and visualized using R software and tidyverse package (81) for statistical analysis.

### JBIR-68 and analog detection in species D isolates

To identify putative analogs of JBIR-68, crude extracts were prepared from five species D isolates. Each isolate was cultured in 20 mL of yeast peptone mannitol (YPM) medium supplemented with HP-20 resin at 28 °C for 10 days. Following cultivation, crude extracts were obtained and subjected to fractionation using ENVplus column chromatography. Sequential elution with increasing concentrations of methanol (25%, 50%, 75%, and 100%) yielded four fractions (F1–F4). LC-MS/MS-based metabolomic analysis was performed on the 100% methanol-eluted fraction (F4) as described above. The acquired metabolomics data was processed using GNPS molecular networking (51). LCMS/MS data for this analysis were also deposited in the MassIVE public repository (MSV000100884).

## Notes

### Competing Interest Statement

The authors have declared no competing interest.

## References

1. Pham JV, Yilma MA, Feliz A, Majid MT, Maffetone N, Walker JR, et al. A Review of the Microbial Production of Bioactive Natural Products and Biologics. Front Microbiol. 2019 Jun 20;10:1404.

2. Lacey HJ, Rutledge PJ. Recently Discovered Secondary Metabolites from Streptomyces Species. Molecules. 2022 Jan 28;27(3):887.

3. Donald L, Pipite A, Subramani R, Owen J, Keyzers RA, Taufa T. Streptomyces: Still the Biggest Producer of New Natural Secondary Metabolites, a Current Perspective. Microbiology Research. 2022 Jul 1;13(3):418–65.

4. Antido JWA, Climacosa FMM. Enhanced Isolation of Streptomyces from Different Soil Habitats in Calamba City, Laguna, Philippines using a Modified Integrated Approach. International Journal of Microbiology. 2022 Oct 26;2022:1–7.

5. Deng Z, Yang W, Lin T, Wang Y, Hua X, Jiang X, et al. Multidimensional insights into the biodiversity of *Streptomyces* in soils of China: a pilot study. Microbiol Spectr. 2025 May 6;13(5):e01692–24.

6. Viaene T, Langendries S, Beirinckx S, Maes M, Goormachtig S. *Streptomyces* as a plant’s best friend? FEMS Microbiology Ecology. 2016 Aug;92(8):fiw119.

7. Liu H, Li J, Singh BK. Harnessing co-evolutionary interactions between plants and Streptomyces to combat drought stress. Nat Plants. 2024 Jul 24;10(8):1159–71.

8. Berasategui A, Shukla S, Salem H, Kaltenpoth M. Potential applications of insect symbionts in biotechnology. Appl Microbiol Biotechnol. 2016 Feb;100(4):1567–77.

9. Chevrette MG, Carlson CM, Ortega HE, Thomas C, Ananiev GE, Barns KJ, et al. The antimicrobial potential of Streptomyces from insect microbiomes. Nat Commun. 2019 Jan 31;10(1):516.

10. Scott JJ, Oh DC, Yuceer MC, Klepzig KD, Clardy J, Currie CR. Bacterial Protection of Beetle-Fungus Mutualism. Science. 2008 Oct 3;322(5898):63–63.

11. Kaltenpoth M. Actinobacteria as mutualists: general healthcare for insects? Trends in Microbiology. 2009 Dec;17(12):529–35.

12. Book AJ, Lewin GR, McDonald BR, Takasuka TE, Doering DT, Adams AS, et al. Cellulolytic Streptomyces Strains Associated with Herbivorous Insects Share a Phylogenetically Linked Capacity To Degrade Lignocellulose. Appl Environ Microbiol. 2014 Aug;80(15):4692–701.

13. Van Arnam EB, Currie CR, Clardy J. Defense contracts: molecular protection in insect-microbe symbioses. Chem Soc Rev. 2018;47(5):1638–51.

14. Van Moll L, De Smet J, Cos P, Van Campenhout L. Microbial symbionts of insects as a source of new antimicrobials: a review. Critical Reviews in Microbiology. 2021 Sep 3;47(5):562–79.

15. Grundmann CO, Guzman J, Vilcinskas A, Pupo MT. The insect microbiome is a vast source of bioactive small molecules. Nat Prod Rep. 2024;41(6):935–67.

16. Carter DO, Yellowlees D, Tibbett M. Cadaver decomposition in terrestrial ecosystems. Naturwissenschaften. 2006 Dec 13;94(1):12–24.

17. Parmenter RR, MacMahon JA. Carrion decomposition and nutrient cycling in a semiarid shrub–steppe ecosystem. Ecological Monographs. 2009 Nov;79(4):637–61.

18. Metcalf JL, Xu ZZ, Weiss S, Lax S, Van Treuren W, Hyde ER, et al. Microbial community assembly and metabolic function during mammalian corpse decomposition. Science. 2016 Jan 8;351(6269):158–62.

19. Wilson EE, Wolkovich EM. Scavenging: how carnivores and carrion structure communities. Trends in Ecology & Evolution. 2011 Mar;26(3):129–35.

20. Metcalf JL, Wegener Parfrey L, Gonzalez A, Lauber CL, Knights D, Ackermann G, et al. A microbial clock provides an accurate estimate of the postmortem interval in a mouse model system. eLife. 2013 Oct 15;2.

21. Guo J, Fu X, Liao H, Hu Z, Long L, Yan W, et al. Potential use of bacterial community succession for estimating post-mortem interval as revealed by high-throughput sequencing. Sci Rep. 2016 Apr 7;6(1).

22. Hyde ER, Haarmann DP, Lynne AM, Bucheli SR, Petrosino JF. The Living Dead: Bacterial Community Structure of a Cadaver at the Onset and End of the Bloat Stage of Decomposition. PLoS ONE. 2013 Oct 30;8(10):e77733.

23. Dekeirsschieter J, Verheggen FJ, Gohy M, Hubrecht F, Bourguignon L, Lognay G, et al. Cadaveric volatile organic compounds released by decaying pig carcasses (Sus domesticus L.) in different biotopes. Forensic Science International. 2009 Aug;189(1–3):46–53.

24. Jordan H, Tomberlin J. Abiotic and Biotic Factors Regulating Inter-Kingdom Engagement between Insects and Microbe Activity on Vertebrate Remains. Insects. 2017 May 24;8(2):54.

25. Thümmel L, Lutz L, Geissenberger J, Pittner S, Heimer J, Amendt J. Decomposition and insect succession of pig cadavers in tents versus outdoors – A preliminary study. Forensic Science International. 2023 May;346:111640.

26. Pechal JL, Crippen TL, Benbow ME, Tarone AM, Dowd S, Tomberlin JK. The potential use of bacterial community succession in forensics as described by high throughput metagenomic sequencing. Int J Legal Med. 2014 Jan;128(1):193–205.

27. Hyde ER, Haarmann DP, Petrosino JF, Lynne AM, Bucheli SR. Initial insights into bacterial succession during human decomposition. Int J Legal Med. 2015 May;129(3):661–71.

28. Takagi M, Motohashi K, Nagai A, Izumikawa M, Tanaka M, Fuse S, et al. Anti-Influenza Virus Compound from *Streptomyces* sp. RI18. Org Lett. 2010 Oct 15;12(20):4664–6.

29. Igarashi Y, Kyoso T, Kim Y, Oikawa T. Simamycin (5′-O-geranyluridine): a new prenylated nucleoside from Streptomyces sp. J Antibiot. 2017 May;70(5):607–10.

30. Parks DH, Chuvochina M, Rinke C, Mussig AJ, Chaumeil PA, Hugenholtz P. GTDB: an ongoing census of bacterial and archaeal diversity through a phylogenetically consistent, rank normalized and complete genome-based taxonomy. Nucleic Acids Research. 2022 Jan 7;50(D1):D785–94.

31. Temkin MI, Carlson CM, Stubbendieck AL, Currie CR, Stubbendieck RM. High Throughput Co-culture Assays for the Investigation of Microbial Interactions. JoVE. 2019 Oct 15;(152):60275.

32. Tenebro CP, Trono DJVL, Vicera CVB, Sabido EM, Ysulat JA, Macaspac AJM, et al. Multiple strain analysis of Streptomyces species from Philippine marine sediments reveals intraspecies heterogeneity in antibiotic activities. Sci Rep. 2021 Sep 2;11(1):17544.

33. Sabido EM, Tenebro CP, Trono DJVL, Vicera CVB, Leonida SFL, Maybay JJWB, et al. Insights into the Variation in Bioactivities of Closely Related Streptomyces Strains from Marine Sediments of the Visayan Sea against ESKAPE and Ovarian Cancer. Marine Drugs. 2021 Jul 31;19(8):441.

34. Blin K, Shaw S, Vader L, Szenei J, Reitz ZL, Augustijn HE, et al. antiSMASH 8.0: extended gene cluster detection capabilities and analyses of chemistry, enzymology, and regulation. Nucleic Acids Research. 2025 Jul 7;53(W1):W32–8.

35. Navarro-Muñoz JC, Selem-Mojica N, Mullowney MW, Kautsar SA, Tryon JH, Parkinson EI, et al. A computational framework to explore large-scale biosynthetic diversity. Nat Chem Biol. 2020 Jan;16(1):60–8.

36. Luo Y, Huang H, Liang J, Wang M, Lu L, Shao Z, et al. Activation and characterization of a cryptic polycyclic tetramate macrolactam biosynthetic gene cluster. Nat Commun. 2013 Dec 5;4(1):2894.

37. Seipke RF, Hutchings MI. The regulation and biosynthesis of antimycins. Beilstein J Org Chem. 2013 Nov 19;9:2556–63.

38. Lechevalier H, Acker RF, Corke CT, Haenseler CM, Waksman SA. Candicidin, A New Antifungal Antibiotic. Mycologia. 1953 Mar;45(2):155–71.

39. Hou L, Huang H, Li H, Wang S, Ju J, Li W. Overexpression of a type III PKS gene affording novel violapyrones with enhanced anti-influenza A virus activity. Microb Cell Fact. 2018 Dec;17(1):61.

40. Pan G, Xu Z, Guo Z, Hindra, Ma M, Yang D, et al. Discovery of the leinamycin family of natural products by mining actinobacterial genomes. Proc Natl Acad Sci USA. 2017 Dec 26;114(52).

41. Martin R, Sierner O, Alvarez MA, Clercq ED, Bailey JE, Minas W. Collinone, a New Recombinant Angular Polyketide Antibiotic Made by an Engineered Streptomyces Strain. J Antibiot. 2001;54(3):239–49.

42. Galmarini OL, Deulofeu V. Curamycin—I. Tetrahedron. 1961 Jan;15(1–4):76–86.

43. Zhao B, Lin X, Lei L, Lamb DC, Kelly SL, Waterman MR, et al. Biosynthesis of the Sesquiterpene Antibiotic Albaflavenone in Streptomyces coelicolor A3(2). Journal of Biological Chemistry. 2008 Mar;283(13):8183–9.

44. Shi J, Zeng YJ, Zhang B, Shao FL, Chen YC, Xu X, et al. Comparative genome mining and heterologous expression of an orphan NRPS gene cluster direct the production of ashimides. Chem Sci. 2019;10(10):3042–8.

45. Bihlmaier C, Welle E, Hofmann C, Welzel K, Vente A, Breitling E, et al. Biosynthetic Gene Cluster for the Polyenoyltetramic Acid α-Lipomycin. Antimicrob Agents Chemother. 2006 Jun;50(6):2113–21.

46. Bikash B, Vilja S, Mitchell L, Keith Y, Mikael I, Mikko MK, et al. Differential regulation of undecylprodigiosin biosynthesis in the yeast-scavenging *Streptomyces* strain MBK6. FEMS Microbiology Letters. 2021 May 6;368(8):fnab044.

47. Thanapipatsiri A, Gomez-Escribano JP, Song L, Bibb MJ, Al-Bassam M, Chandra G, et al. Discovery of Unusual Biaryl Polyketides by Activation of a Silent *Streptomyces venezuelae* Biosynthetic Gene Cluster. ChemBioChem. 2016 Nov 17;17(22):2189–98.

48. Becerril A, Álvarez S, Braña AF, Rico S, Díaz M, Santamaría RI, et al. Uncovering production of specialized metabolites by Streptomyces argillaceus: Activation of cryptic biosynthesis gene clusters using nutritional and genetic approaches. Virolle MJ, editor. PLoS ONE. 2018 May 24;13(5):e0198145.

49. Wernke KM, Xue M, Tirla A, Kim CS, Crawford JM, Herzon SB. Structure and bioactivity of colibactin. Bioorganic & Medicinal Chemistry Letters. 2020 Aug;30(15):127280.

50. Inahashi Y, Zhou S, Bibb MJ, Song L, Al-Bassam MM, Bibb MJ, et al. Watasemycin biosynthesis in Streptomyces venezuelae: thiazoline C-methylation by a type B radical-SAM methylase homologue. Chem Sci. 2017;8(4):2823–31.

51. Wang M, Carver JJ, Phelan VV, Sanchez LM, Garg N, Peng Y, et al. Sharing and community curation of mass spectrometry data with Global Natural Products Social Molecular Networking. Nat Biotechnol. 2016 Aug;34(8):828–37.

52. Mohimani H, Gurevich A, Shlemov A, Mikheenko A, Korobeynikov A, Cao L, et al. Dereplication of microbial metabolites through database search of mass spectra. Nat Commun. 2018 Oct 2;9(1):4035.

53. Birch AJ, Cameron DW, Harada Y, Rickards RW. 187. The structure of the antimycin-a complex. J Chem Soc. 1961;889.

54. Takada K, Ninomiya A, Naruse M, Sun Y, Miyazaki M, Nogi Y, et al. Surugamides A–E, Cyclic Octapeptides with Four D-Amino Acid Residues, from a Marine Streptomyces sp.: LC–MS-Aided Inspection of Partial Hydrolysates for the Distinction of D- and L-Amino Acid Residues in the Sequence. J Org Chem. 2013 Jul 5;78(13):6746–50.

55. Schumacher RW, Talmage SC, Miller SA, Sarris KE, Davidson BS, Goldberg A. Isolation and Structure Determination of an Antimicrobial Ester from a Marine Sediment-Derived Bacterium. J Nat Prod. 2003 Sep 1;66(9):1291–3.

56. Vicente CM, Thibessard A, Lorenzi JN, Benhadj M, Hôtel L, Gacemi-Kirane D, et al. Comparative Genomics among Closely Related Streptomyces Strains Revealed Specialized Metabolite Biosynthetic Gene Cluster Diversity. Antibiotics. 2018 Oct 2;7(4):86.

57. Sangwan NK, Verma BS, Verma KK, Dhindsa KS. Nematicidal activity of some essential plant oils. Pestic Sci. 1990 Jan;28(3):331–5.

58. Echeverrigaray S, Zacaria J, Beltrão R. Nematicidal Activity of Monoterpenoids Against the Root-Knot Nematode *Meloidogyne incognita*. Phytopathology®. 2010 Feb;100(2):199–203.

59. Laquale S, Candido V, Avato P, Argentieri MP, D’Addabbo T. Essential oils as soil biofumigants for the control of the root-knot nematode *Meloidogyne incognita* on tomato: Essential oils as soil biofumigants. Ann Appl Biol. 2015 Sep;167(2):217–24.

60. Panda SK, Daemen M, Sahoo G, Luyten W. Essential Oils as Novel Anthelmintic Drug Candidates. Molecules. 2022 Nov 29;27(23):8327.

61. Mathison BA, Couturier MR, Pritt BS. Diagnostic Identification and Differentiation of Microfilariae. Kraft CS, editor. J Clin Microbiol. 2019 Oct;57(10):e00706–19.

62. Hayakawa M, Nonomura H. Humic acid-vitamin agar, a new medium for the selective isolation of soil actinomycetes. Journal of Fermentation Technology. 1987 Jan;65(5):501–9.

63. Hanshew AS, McDonald BR, Díaz Díaz C, Djiéto-Lordon C, Blatrix R, Currie CR. Characterization of Actinobacteria Associated with Three Ant–Plant Mutualisms. Microb Ecol. 2015 Jan;69(1):192–203.

64. Liu Y, Schröder J, Schmidt B. Musket: a multistage *k-* mer spectrum-based error corrector for Illumina sequence data. Bioinformatics. 2013 Feb 1;29(3):308–15.

65. Magoč T, Salzberg SL. FLASH: fast length adjustment of short reads to improve genome assemblies. Bioinformatics. 2011 Nov 1;27(21):2957–63.

66. Bankevich A, Nurk S, Antipov D, Gurevich AA, Dvorkin M, Kulikov AS, et al. SPAdes: A New Genome Assembly Algorithm and Its Applications to Single-Cell Sequencing. Journal of Computational Biology. 2012 May;19(5):455–77.

67. Chaumeil PA, Mussig AJ, Hugenholtz P, Parks DH. GTDB-Tk v2: memory friendly classification with the Genome Taxonomy Database. 2022.

68. Salamzade R, Kottapalli A, Kalan LR. skDER and CiDDER: two scalable approaches for microbial genome dereplication. Microbial Genomics. 2025 Jul 10;11(7).

69. Shaw J, Yu YW. Fast and robust metagenomic sequence comparison through sparse chaining with skani. Nat Methods. 2023 Nov;20(11):1661–5.

70. Larralde M. Pyrodigal: Python bindings and interface to Prodigal,an efficient method for gene prediction in prokaryotes. JOSS. 2022 Apr 25;7(72):4296.

71. Lee MD. GToTree: a user-friendly workflow for phylogenomics. Ponty Y, editor. Bioinformatics. 2019 Oct 15;35(20):4162–4.

72. Price MN, Dehal PS, Arkin AP. FastTree 2 – Approximately Maximum-Likelihood Trees for Large Alignments. Poon AFY, editor. PLoS ONE. 2010 Mar 10;5(3):e9490.

73. Menardo F, Loiseau C, Brites D, Coscolla M, Gygli SM, Rutaihwa LK, et al. Treemmer: a tool to reduce large phylogenetic datasets with minimal loss of diversity. BMC Bioinformatics. 2018 Dec;19(1):164.

74. Chklovski A, Parks DH, Woodcroft BJ, Tyson GW. CheckM2: a rapid, scalable and accurate tool for assessing microbial genome quality using machine learning. Nat Methods. 2023 Aug;20(8):1203–12.

75. Letunic I, Bork P. Interactive Tree of Life (iTOL) v6: recent updates to the phylogenetic tree display and annotation tool. Nucleic Acids Research. 2024 Jul 5;52(W1):W78–82.

76. Raivo Kolde. pheatmap: Pretty Heatmaps. 2010. p. 1.0.12.

77. Terlouw BR, Blin K, Navarro-Muñoz JC, Avalon NE, Chevrette MG, Egbert S, et al. MIBiG 3.0: a community-driven effort to annotate experimentally validated biosynthetic gene clusters. Nucleic Acids Research. 2023 Jan 6;51(D1):D603–10.

78. Chambers MC, Maclean B, Burke R, Amodei D, Ruderman DL, Neumann S, et al. A cross-platform toolkit for mass spectrometry and proteomics. Nat Biotechnol. 2012 Oct;30(10):918–20.

79. Michalski ML, Griffiths KG, Williams SA, Kaplan RM, Moorhead AR. The NIH-NIAID Filariasis Research Reagent Resource Center. Knight M, editor. PLoS Negl Trop Dis. 2011 Nov 29;5(11):e1261.

80. Wheeler NJ, Gallo KJ, Rehborg EJG, Ryan KT, Chan JD, Zamanian M. wrmXpress: A modular package for high-throughput image analysis of parasitic and free-living worms. Cotton J, editor. PLoS Negl Trop Dis. 2022 Nov 18;16(11):e0010937.

81. Wickham H, Averick M, Bryan J, Chang W, McGowan L, François R, et al. Welcome to the Tidyverse. JOSS. 2019 Nov 21;4(43):1686.

