## Supplemental data for "Bioactive Natural Products Produced by *Streptomyces* from the Microbiome of Cadaveric Fly Larvae"

### Content

Supplementary Figures and Tables

Figure S1. Phylogenetic tree depicting average nucleotide identity (ANI) of *Streptomyces* isolates recovered from cadaveric fly larvae

Figure S2. Taxonomic classification of *Streptomyces* isolates recovered from cadaveric fly larvae.

Figure S3. GNPS rendered molecular network of crude metabolic extracts generated from *Streptomyces* isolates

Figure S4. MS1 for detected antimycins with commercially available antimycin standard

Figure S5. MS1 for detected surugamides A & B

Figure S6. MS1 for detected surugamide G

Figure S7. MS1 for detected surugamide H

Figure S8. MS1 for detected Macrotetrolide

Figure S9. MS1 for detected Bonactin

Figure S10. The 1H NMR spectrum of JBIR-68 (1) in MeOD

Figure S11. The COSY spectrum of JBIR-68 (1) in MeOD

Figure S12. The HSQC spectrum of JBIR-68 (1) in MeOD

Figure S13. The HMBC spectrum of JBIR-68 (1) in MeOD

Figure S14. Comparison of MS2 of purified JBIR-68 and Simamycin

Figure S15. Extracted Ion Chromatogram (EIC) of JBIR-68 and Simamycin obtained from semicrude fraction of SID9885

Figure S16. Extracted Ion Chromatogram (EIC) of JBIR-68 and analogs from semicrude fraction of a species D representative

Table S1. $^{1}H$ and $^{13}C$ NMR Spectroscopic Data for JBIR-68

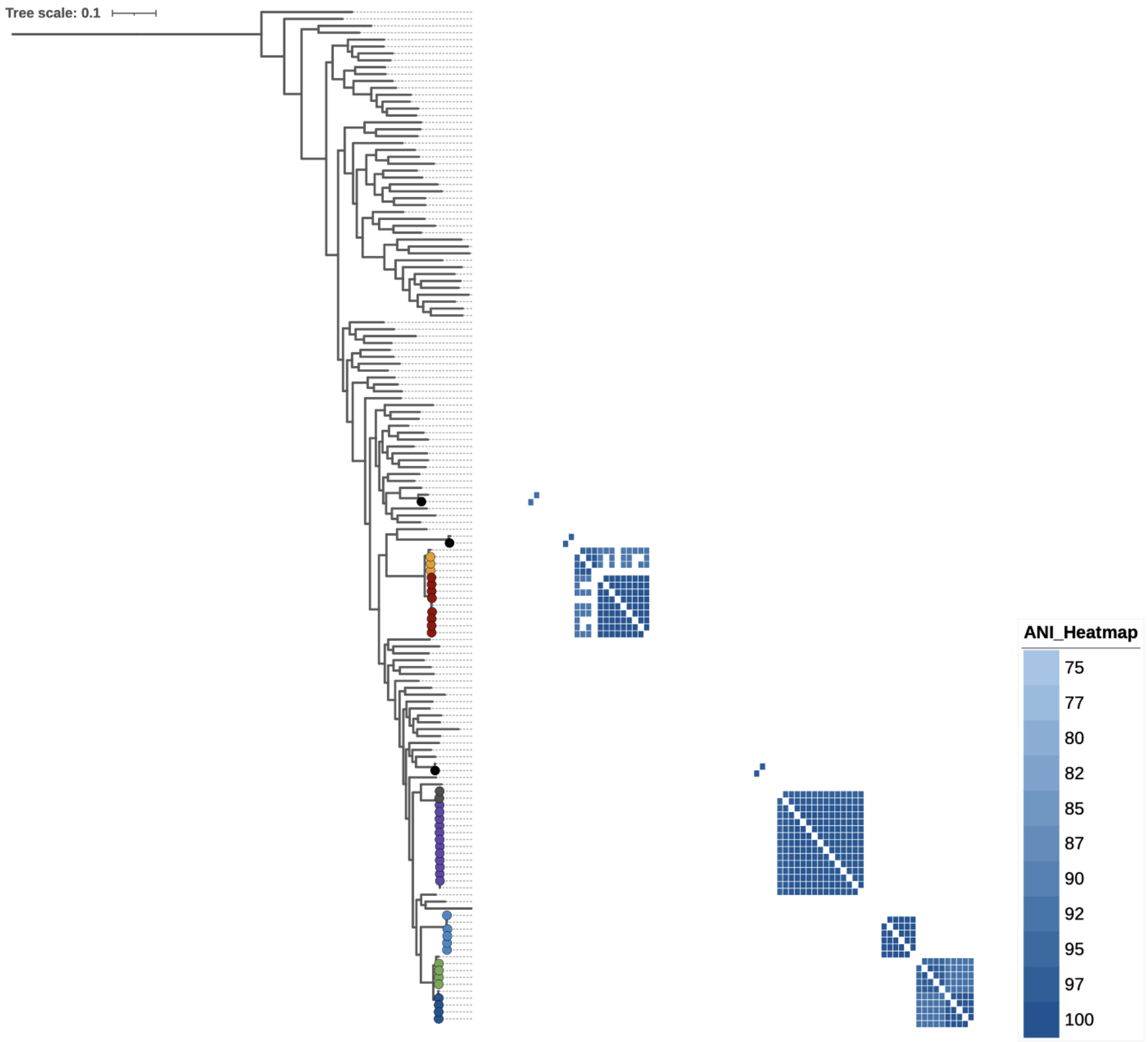


Figure S1. **Phylogenetic tree depicting average nucleotide identity (ANI) of *Streptomyces* isolates recovered from cadaveric fly larvae.** The colored codes indicate *Streptomyces* isolated from the cadaveric fly larvae.

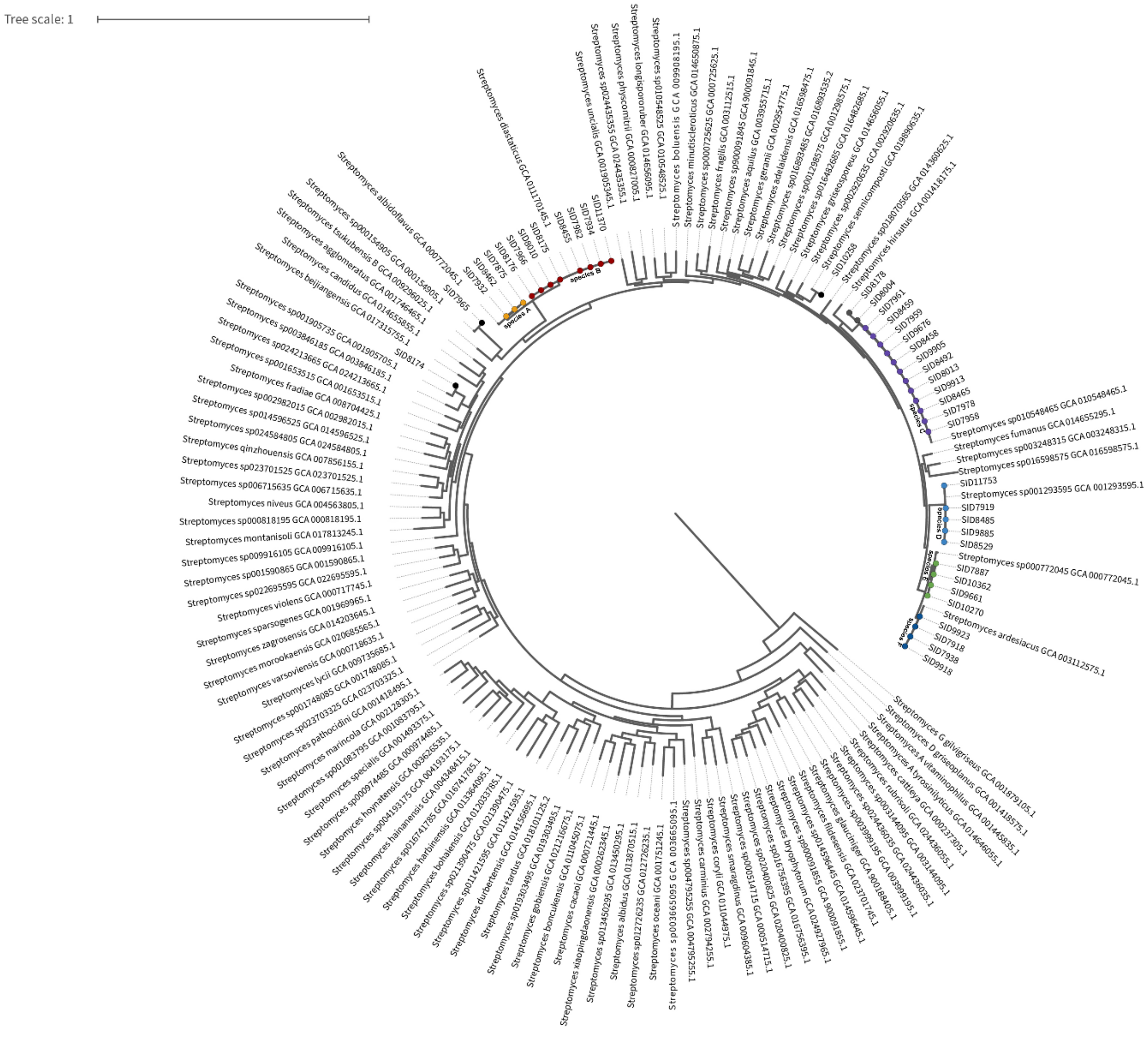


Figure S2. **Taxonomic classification of *Streptomyces* isolates recovered from cadaveric fly larvae.** A genome-based phylogeny was constructed using the 42 *Streptomyces* isolates obtained in this study, six GTDB R214 reference genomes representing the species recovered here, and an additional set of 100 diverse *Streptomyces* genomes to provide broader taxonomic context. The phylogeny was generated from GTDB-Tk concatenated marker genes and visualized in iTOL. Isolates recovered from cadaveric fly larvae are labeled as "SID" and highlighted with colored nodes, each corresponding to one of the six species-level groups (A–F) identified in this study.

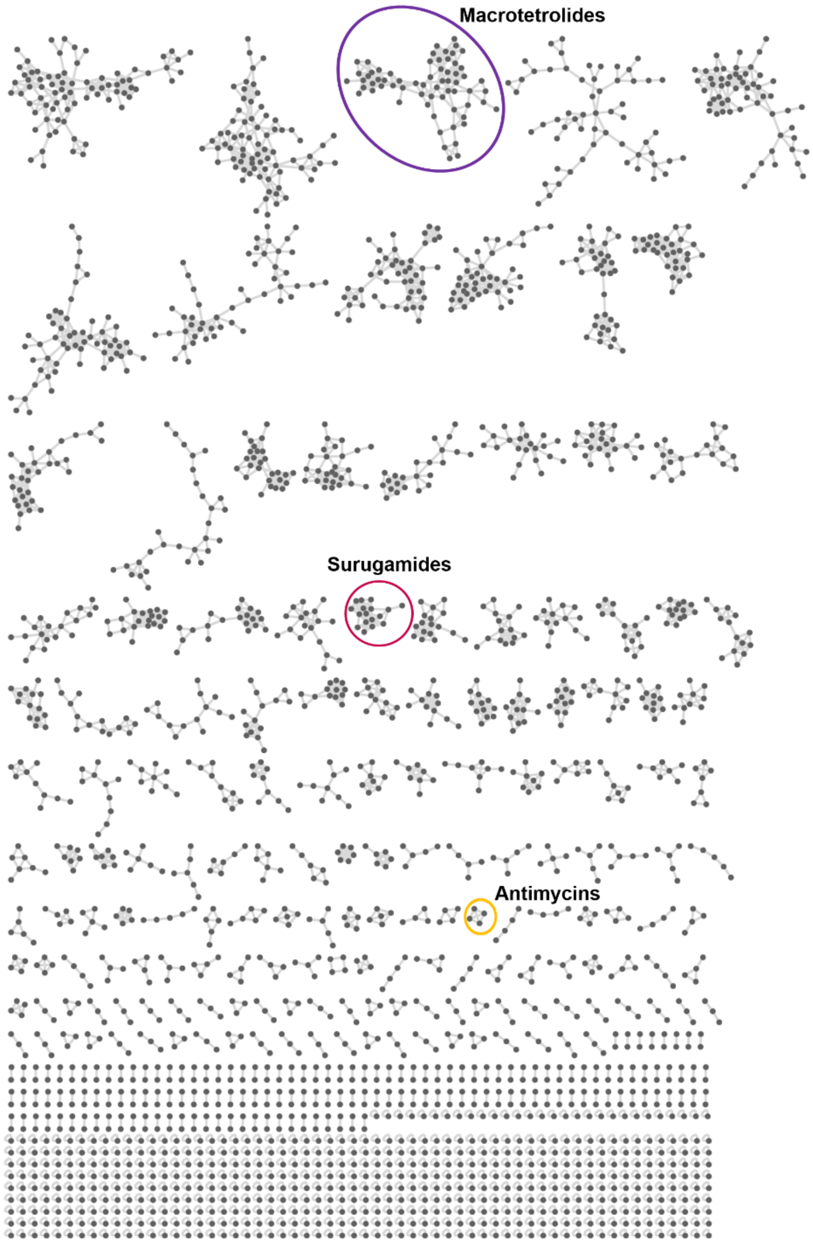


Figure S3. **GNPS rendered molecular network of crude metabolic extracts generated from *Streptomyces* isolates** obtained in this study. The labeled and circled networks were detected as GNPS-library hits. Cytoscape v3.10.1 was utilized to visualize the network.

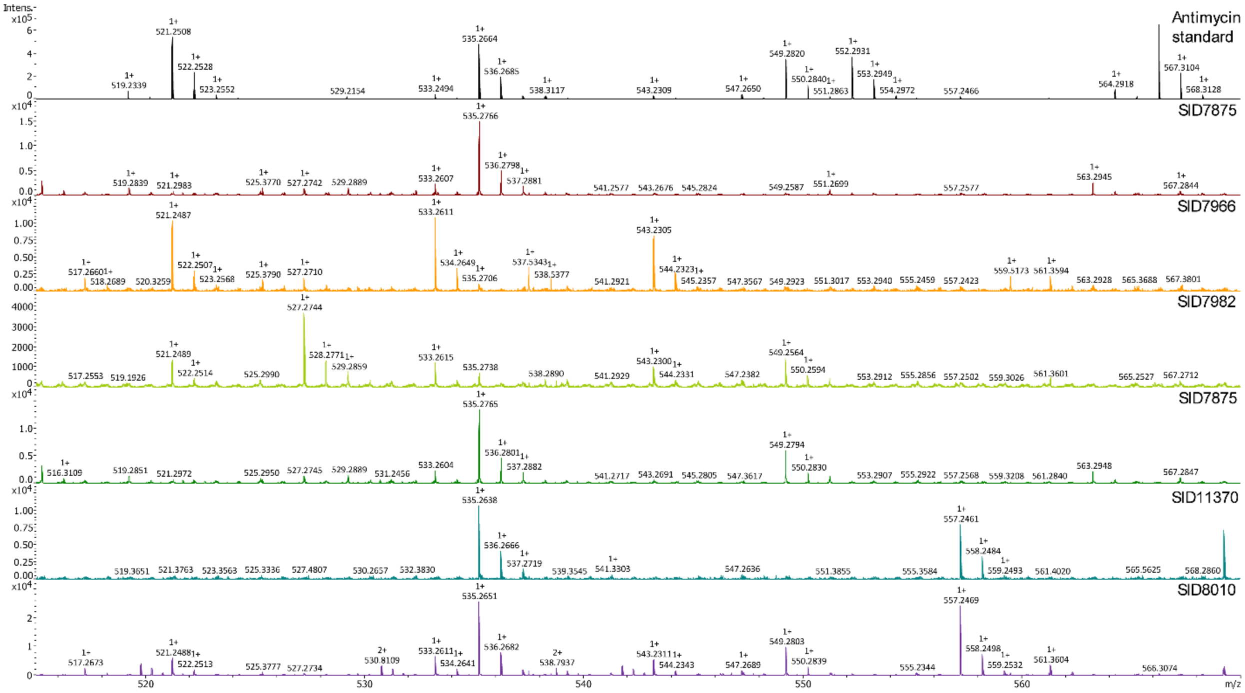


Figure S4. MS1 of detected antimycins in cadaveric fly larvae *Streptomyces* isolates and commercially available antimycin standard.

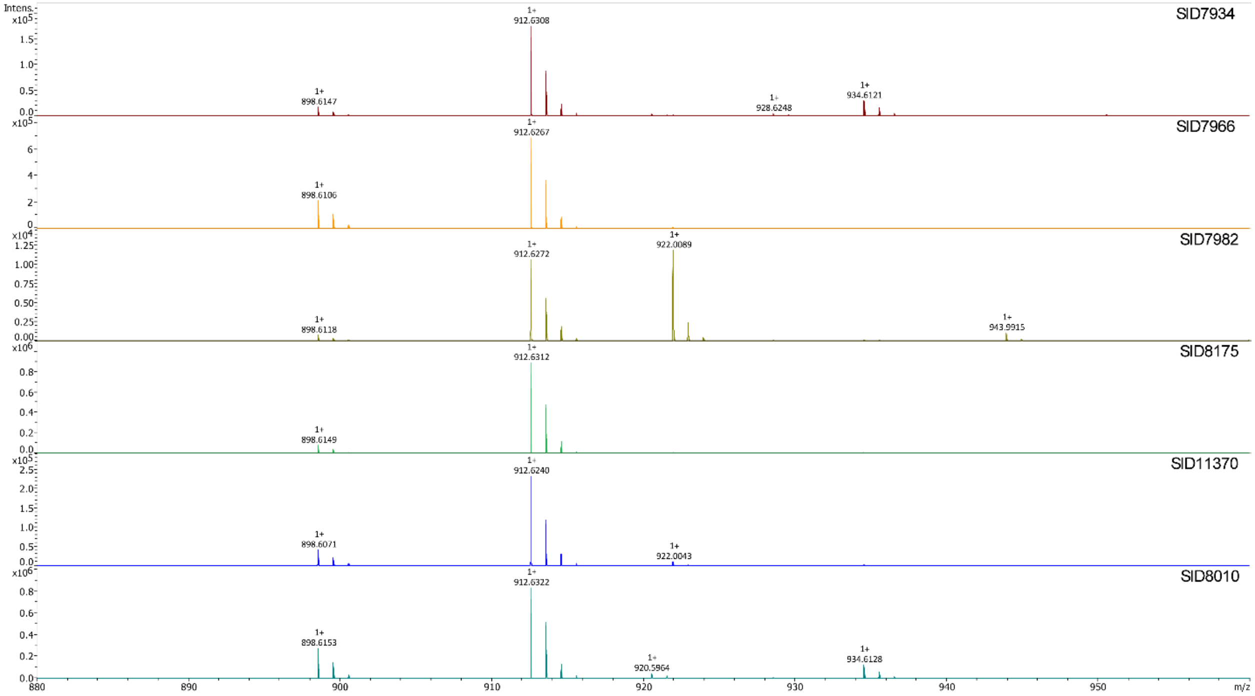


Figure S5. MS1 of detected surugamides A *m/z* 912.024 and Surugamide B *m/z* 898.61 at retention time 9.5-9.7 min in cadaveric fly larvae *Streptomyces* isolates.

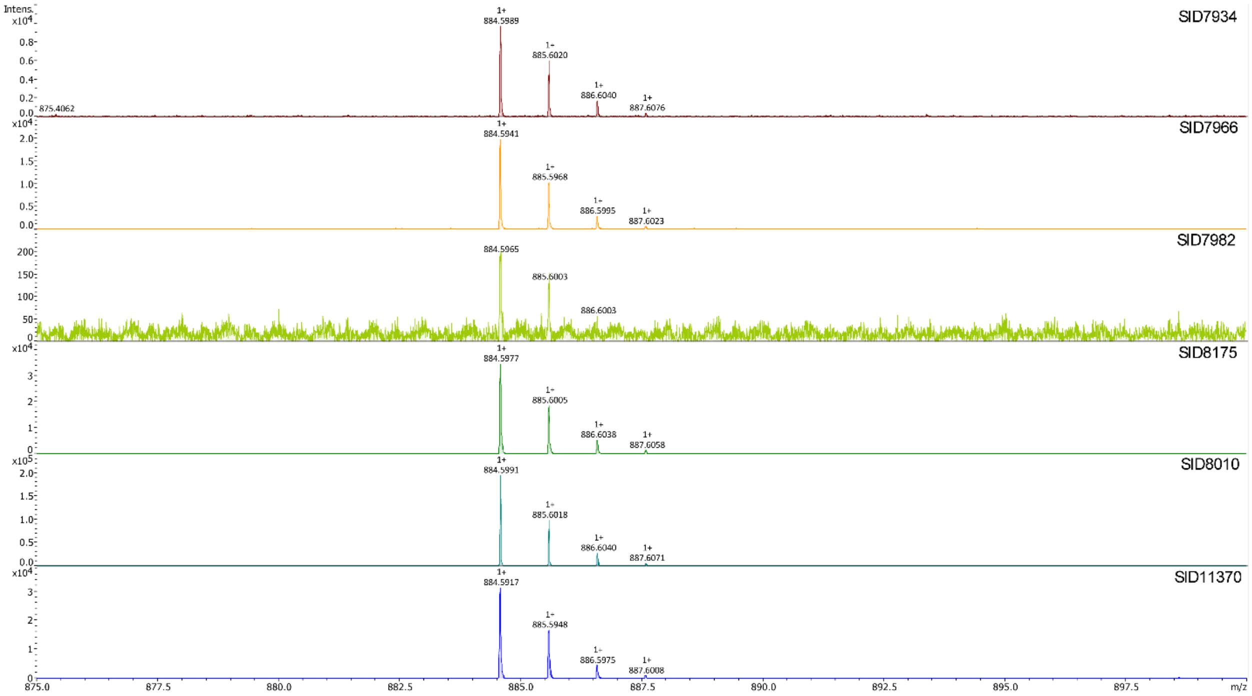


Figure S6. MS1 of detected Surugamides G *m/z* 884.59 at retention time 9.17 min in cadaveric fly larvae *Streptomyces* isolates.

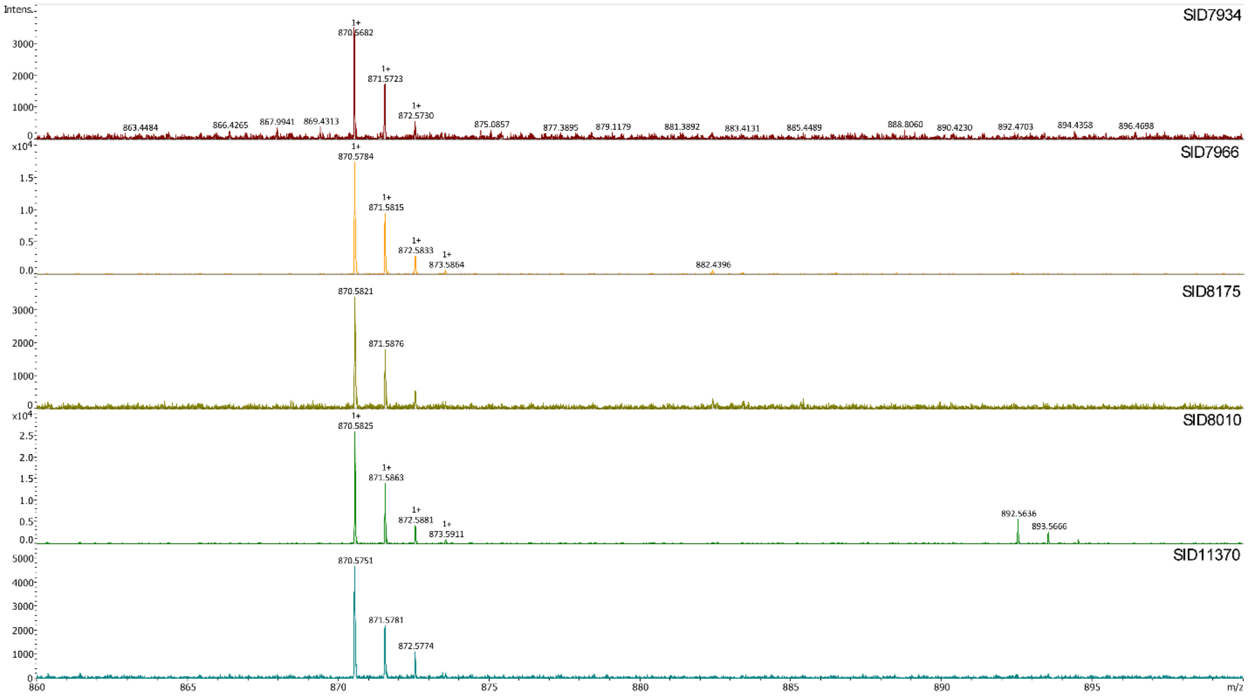


Figure S7. MS1 for detected surugamides H *m/z* 870.58 at retention time 9.11 min in cadaveric fly larvae *Streptomyces* isolates.

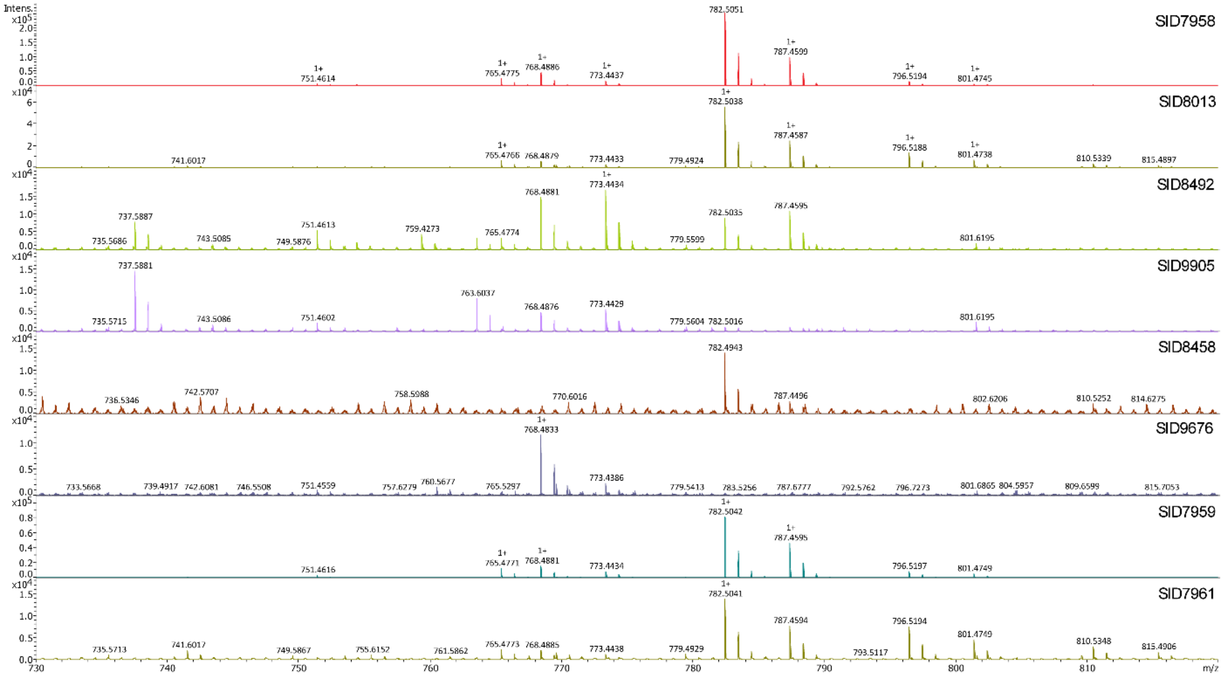


Figure S8. MS1 for detected Macrotetrolide at retention time 12.28-13.30 min in cadaveric fly larvae *Streptomyces* isolates.

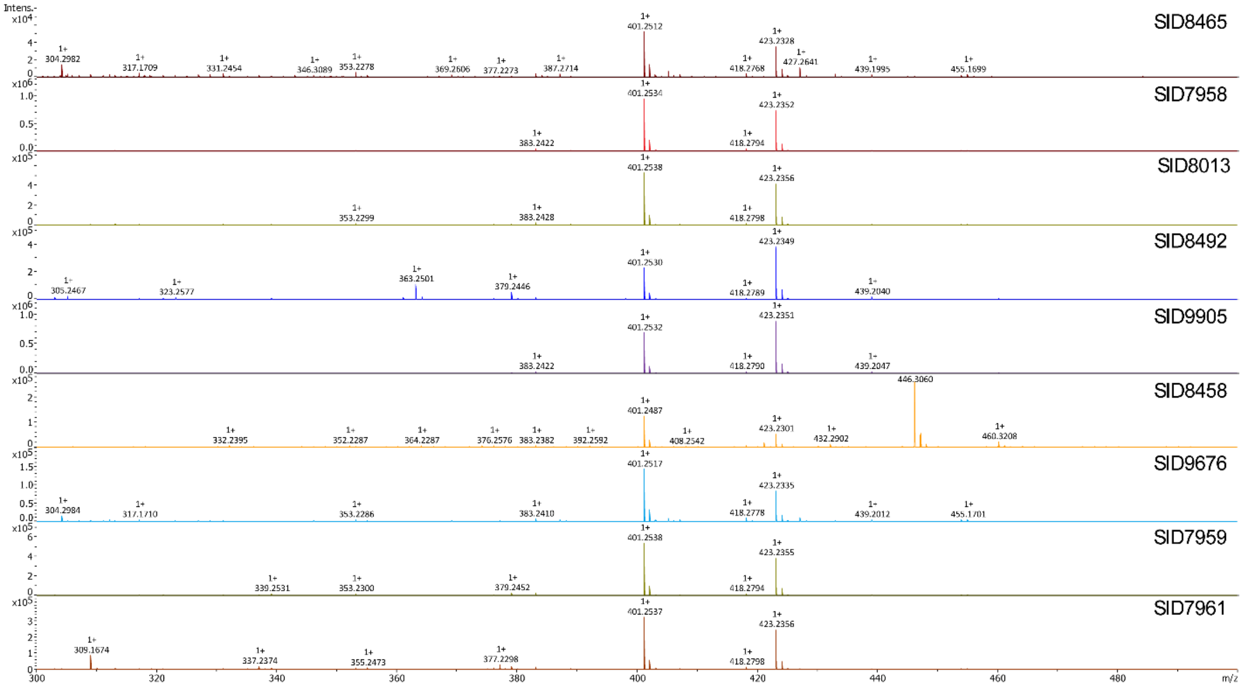


Figure S9. MS1 for detected Bonactin m/z 423.234 at retention time 12.41 min.

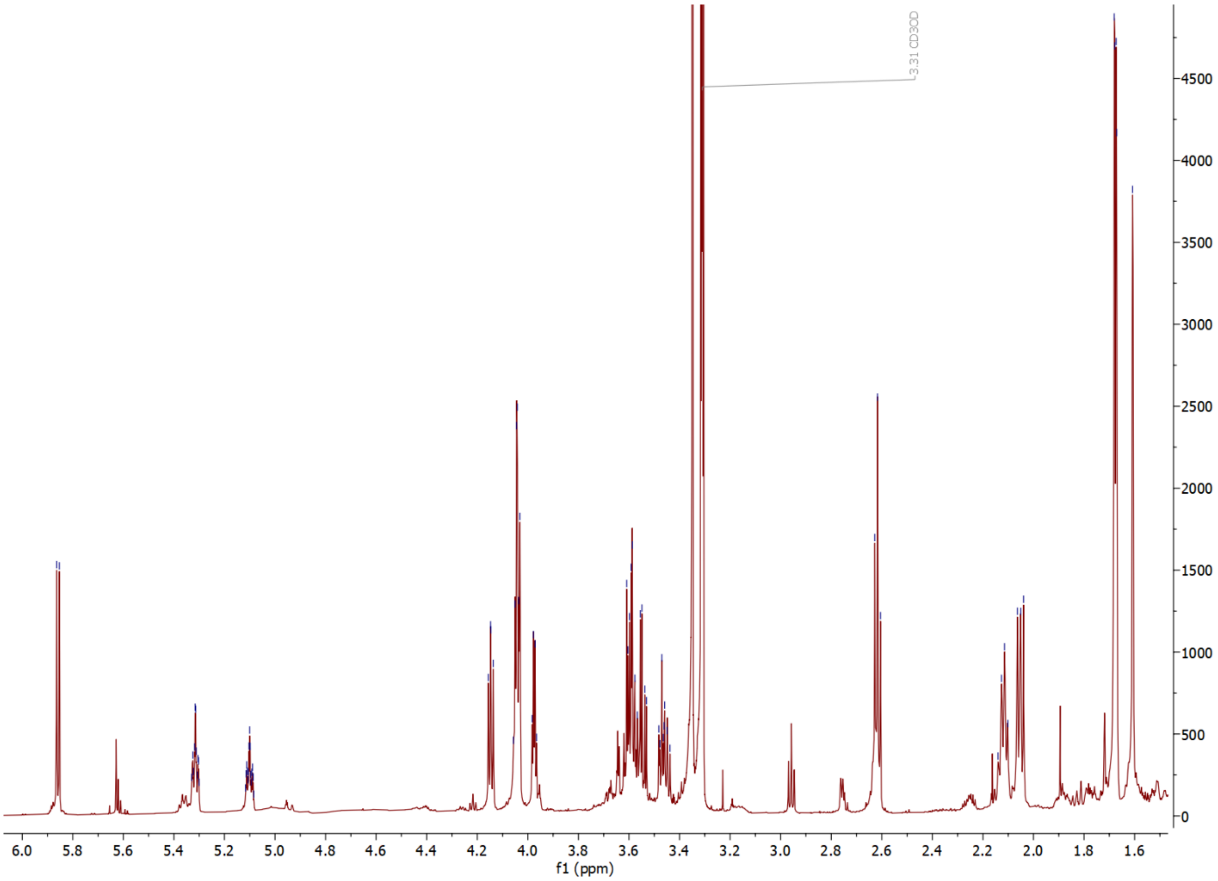


Figure S10. The 1H NMR spectrum (600 MHz) of JBIR-68 (1) in MeOD

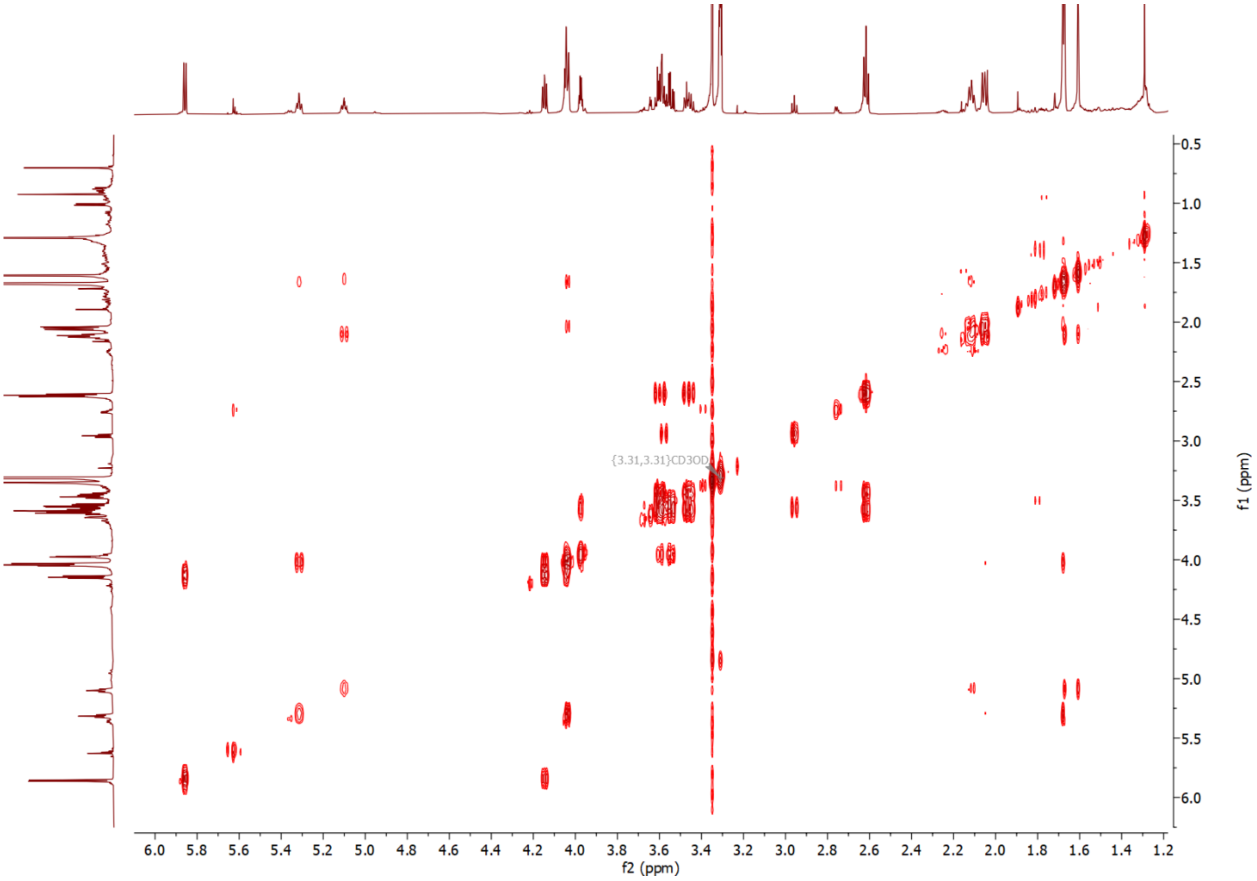


Figure S11. The COSY spectrum (600 MHz) of JBIR-68 (1) in MeOD

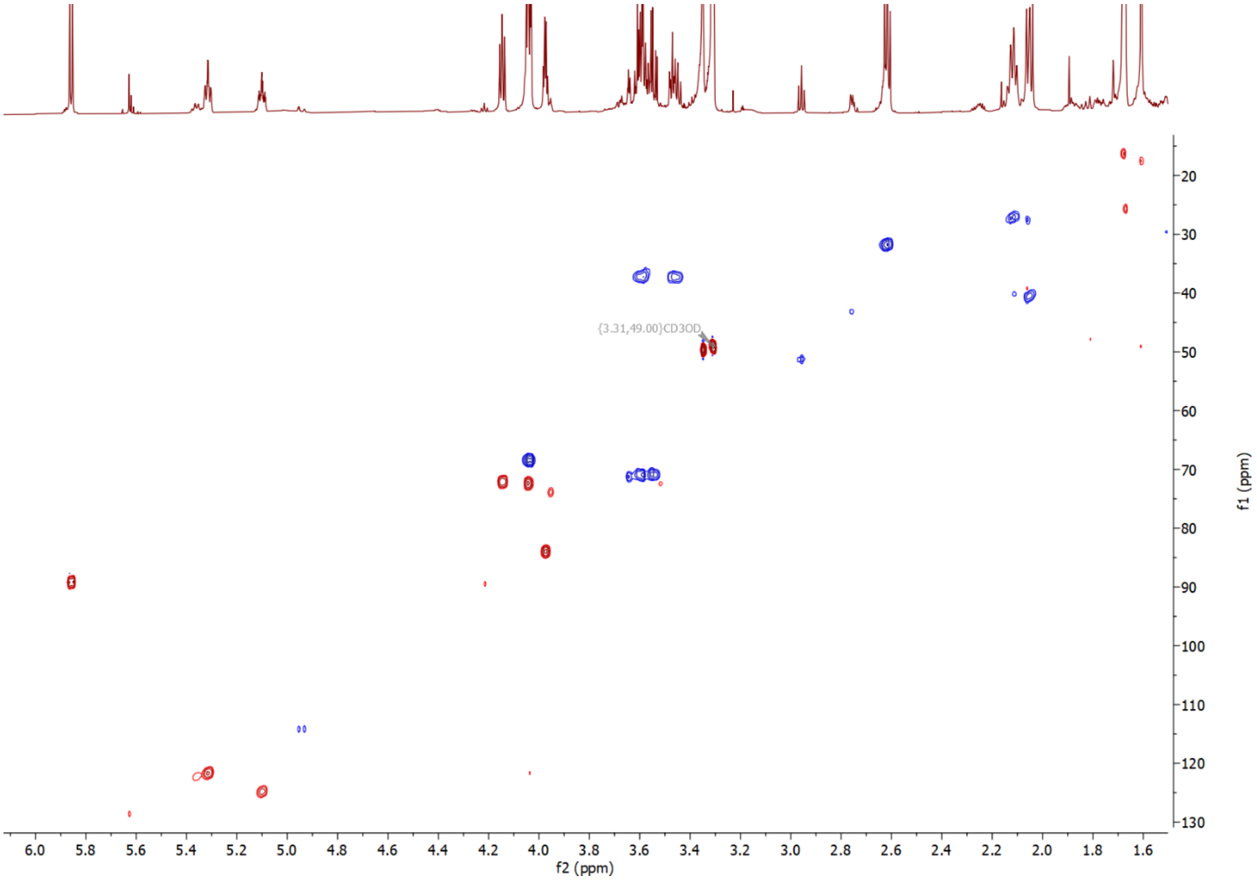


Figure S12. The HSQC spectrum (600 MHz) of JBIR-68 (1) in MeOD

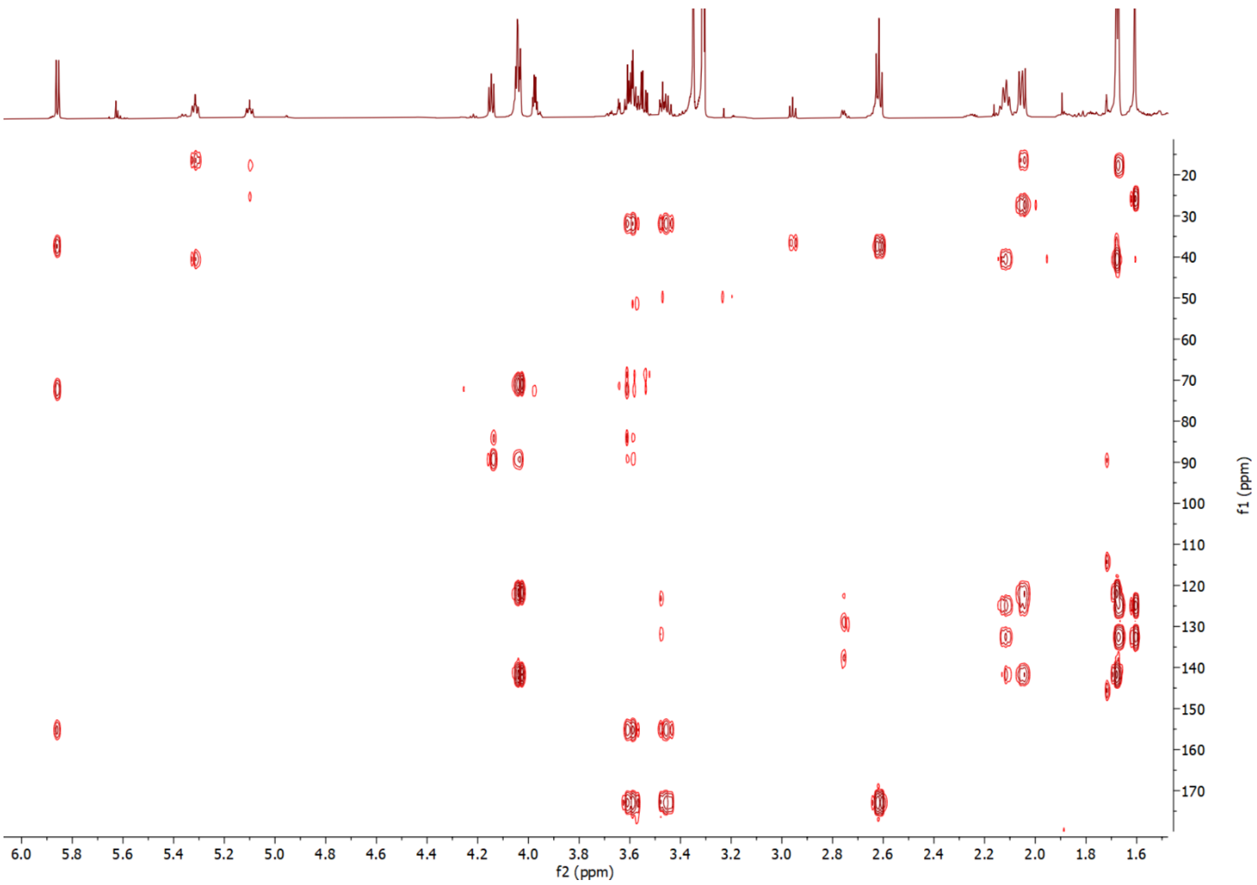


Figure S13. The HMBC spectrum (600 MHz) of JBIR-68 (1) in MeOD

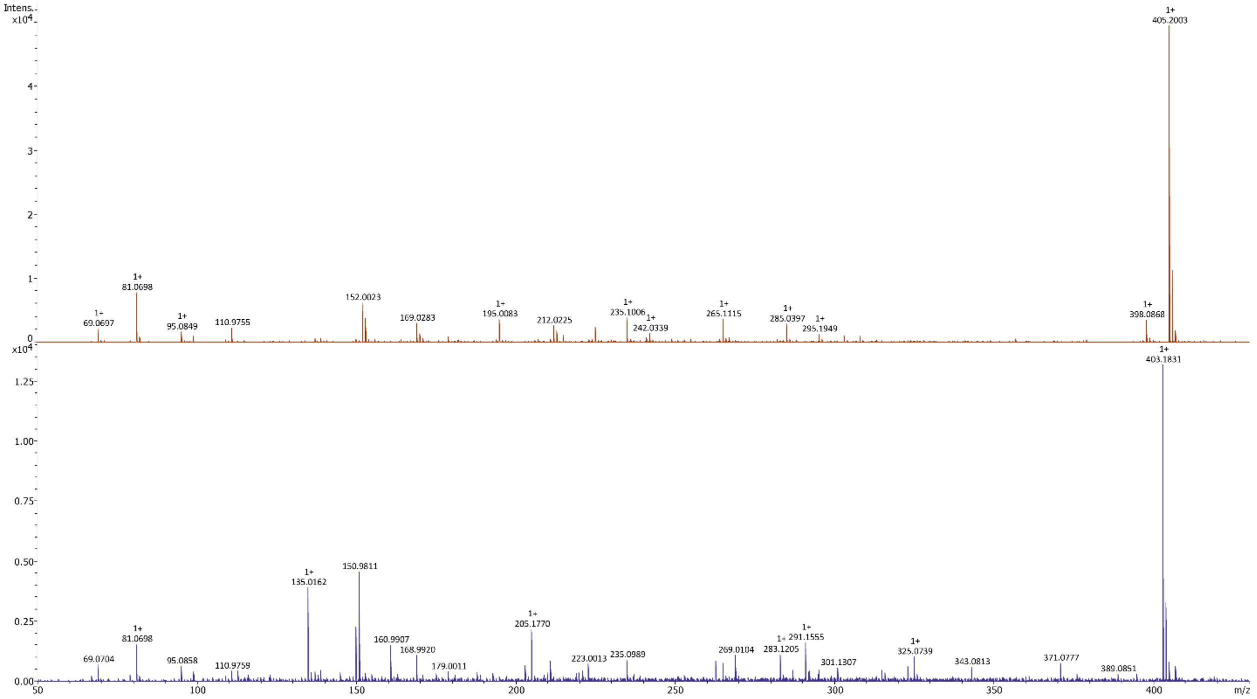


Figure S14. **Comparison of MS2 of purified JBIR-68 and Simamycin.** MS2 spectra of JBIR-68 ($[M+Na]^+$, 405.200) detected and Simamycin ($[M+Na]^+$, 403.183). Several identical fragment ions between JBIR-68 and Simamycin are identifiable. The spectra are obtained from purified compounds.

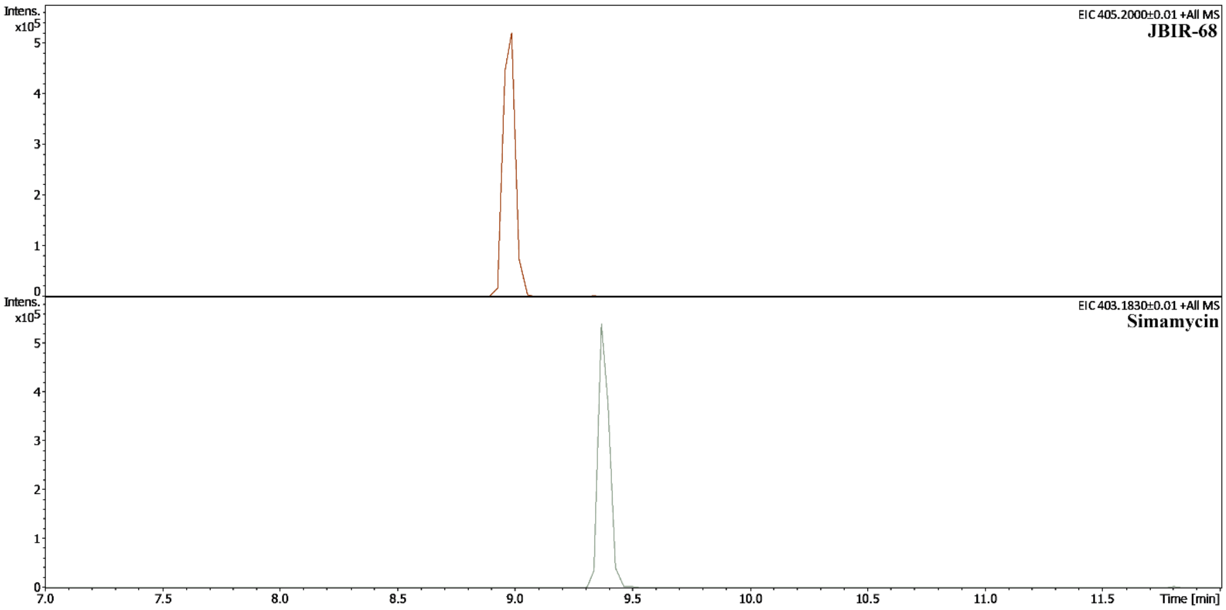


Figure S15. **Extracted Ion Chromatogram (EIC) of JBIR-68 and Simamycin obtained from semicrude fraction of SID9885.** A representative EIC of JBIR-68 detected at retention time of 8.99 min and Simamycin at retention time of 9.37 min. The EIC is rendered from the semicrude fraction of SID9885.

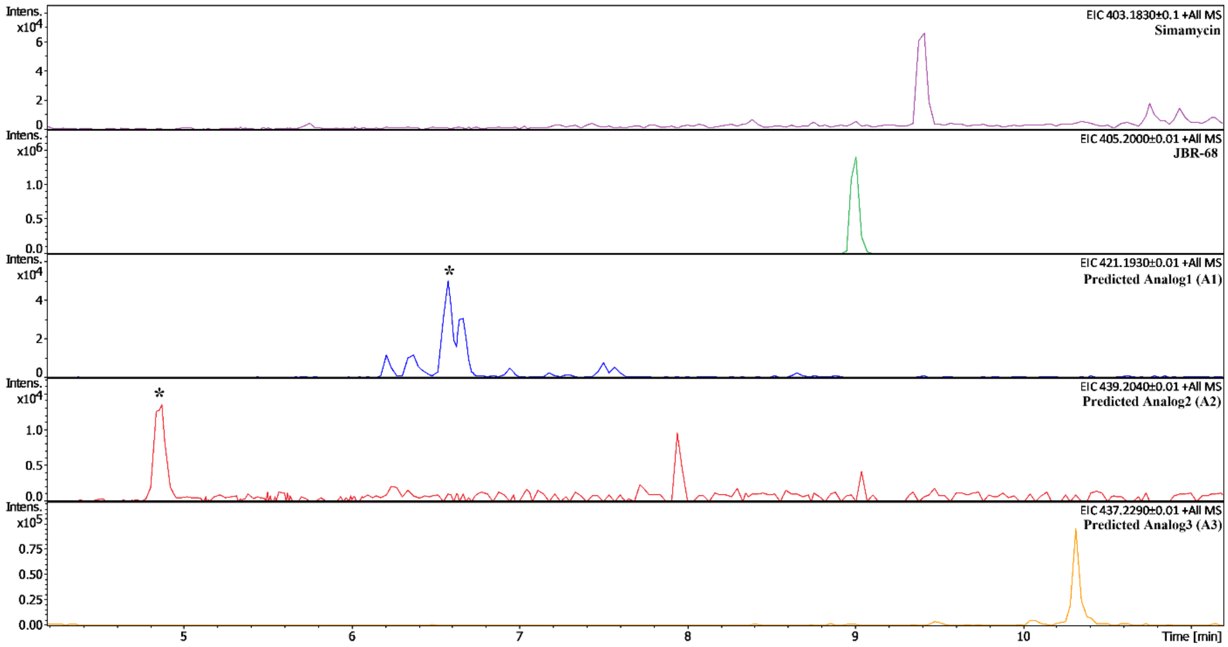


Figure S16. **Extracted Ion Chromatogram (EIC) of JBIR-68 and analogs from semicrude fraction of a species D representative.** EIC of Simamycin, JBIR-68, Predicted Analog 1 (A1), Predicted Analog 2 (A2), and Predicted Analog 3 (A3) from SID8464. EIC indicate different retention times for all these compounds. * Indicates the peak with the detected mass feature.

Table S1. $^{1}H$ (600 MHz) and $^{13}C$ (125 MHz) NMR Spectroscopic Data for JBIR-68 (1) in methanol-$d_4$ ($\delta$ in ppm, $J$ values in Hz)

| Position | 1 | |
|---|---|---|
| | $\delta_C$ | $\delta_H$ ($J$ in Hz) |
| 1 | 68.5 | 4.03[a] |
| 2 | 121.8 | 5.31, m |
| 3 | 141.8 | |
| 4 | 40.6 | 2.05, m |
| 5 | 27.3 | 2.11, m |
| 6 | 124.9 | 5.10, tp |
| 7 | 132.6 | |
| 8 | 25.8 | 1.67, s |
| 9 | 17.7 | 1.61, s |
| 10 | 16.4 | 1.68, s |
| 2’ | 155.2 | |
| 4’ | 172.9 | |
| 5’ | 31.9 | 2.62, t (6.74) |
| 6a’<br>6b’ | 37.4 | 3.54, dd (3.88, 10.74),<br>3.46 |
| 1” | 89.3 | 5.86, d (6.4) |
| 2” | 72.2 | 4.15, dd (5.4, 6.4) |
| 3” | 72.6 | 4.04[a] |
| 4” | 84.1 | 3.97, q (3.12, 3.12, 3.18) |
| 5” | 71.0 | 3.60, m |
